# Co-fluctuations of neural activity define intra-V1 networks related to perceptual organization

**DOI:** 10.1101/2022.08.22.504869

**Authors:** Mitchell Valdes-Sosa, Marlis Ontivero-Ortega, Jorge Iglesias-Fuster, Agustin Lage-Castellanos, Lidice Galan-Garcia, Pedro Valdes-Sosa

## Abstract

Using functional resonance imaging (fMRI), we studied the relationship between perceptual organization and network topology within the primary visual cortex (V1). Twenty-six humans (male and female) were recorded during active observation of two Global and two Local Navon letters. Correlations between fMRI fluctuations from different V1 sites were measured (after removing stimulus-evoked signals) in windows specific to each condition. Intra-V1, like brain-wide networks, presented an overall decrease of correlations during stimulation compared to baseline and increased statistical dimensionality. Massive edgewise testing and network based-statistics (both corrected by FDR) identified differences between conditions of connection strengths that were mapped to the visual field. Global letters elicited long links mainly connecting V1 sites mapping the lower left/right visual quadrants. Shorter links were associated with Local letters, primarily mapped within the lower-left visual quadrant. Frequently link lengths exceeded V1 population receptive field sizes. These connections were not observed in the time-locked (feedforward) responses shared across participants. Thus, these networks reflect activity idiosyncratic to each participant, possibly generated by interactions within or feedback to V1. Perception would sculpt V1 connectivity, with specific increases in link strengths (in a background of decreases). These findings could help shed light on V1 as a “cognitive blackboard”.

## Introduction

The primary visual cortex (V1) is the main recipient of visual input from the retina, and the initial station of specialized -feedforward- (Lamme & Roelfsema, 2000) routes enlisting higher-order visual areas (Goodale & Milner, 2018; Haak & Beckmann, 2016). Animal studies indicate that V1 neurons have small receptive fields (RF) narrowly tuned for different stimulus features (van Essen et al., 1984). The largest sizes of human V1 population RFs (pRFs) (Wandell & Winawer, 2015), estimated with functional magnetic resonance imaging (fMRI), do not exceed 5° of visual angle in V1. These results indicate that V1 is a bank of local feature detectors that provide a basic description of visual scenes (DeValois & DeValois, 1991).

Visual perceptual organization is a critical stage that groups different stimulus elements, separates figure from background, establishes whole/parts relations, and enables object-based attention (Kimchi et al., 2016; Pomerantz & Portillo, 2015; Wagemans et al., 2012). These processes frequently encompass regions in the visual field (VF) spanning distances bigger than pRF sizes in V1. Thus, perceptual organization needs information integration across sizeable visual field sectors, usually thought to be achieved in higher-order visual areas with larger RFs (Klink et al., 2021; Wandell & Winawer, 2015).

Interestingly, V1 contains information about global stimulus properties that extend across distances bigger than its small RFs. Multivariate pattern analysis (MVPA) of human fMRI activity provides evidence for this. Scene identity can be decoded within regions of V1 that map large but occluded parts of the VF (Smith & Muckli, 2010). Object shapes presented in the VF periphery can be decoded from the (unstimulated) foveal V1 (Williams et al., 2008). MVPA correctly predicts visual working-memory content from V1 activity without exogenous stimulation (Harrison & Tong, 2009), reflecting purely top-down information. This enriched V1 activity, not explainable by its classical RFs, could play an essential role in processes such as attention, working memory, and imagery, possibly acting like a cognitive blackboard (Roelfsema & de Lange, 2016).

Interactions between neuronal populations are needed to represent large-scale -global-properties in V1. Lateral connections between V1 neurons having similar edge-orientation could play a role, facilitating the detection of extended contours (Chen et al., 2014; Kapadia et al., 1995; Liang et al., 2017; Schwarz & Bolz, 1991). But input from higher-order visual areas (Roelfsema & de Lange, 2016) is required to make more complex features (e.g., object form or scene identity) and broader spatial coverage available to V1. Massive fiber connections from the visual higher-order regions to V1 could provide an anatomical basis for this feedback (Markov et al., 2011; Markov et al., 2014).

To understand perceptual organization, we must measure cooperation between neurons scattered within V1, higher-order visual cortices, and beyond. Doing so is a daunting task. Luckily, this cooperation may imply the synchronization of neural activity (Singer, 2021), which could perhaps be measured in V1. Brain sites with synchronized slow fMRI signals are parts of the same large-scale brain circuits (Bijsterbosch et al., 2017, Bijsterbosch et al., 2020).

Encouragingly for our purposes, fine-grained patterns of visual neural circuitry are also reflected by this type of synchronization. Connective receptive field (cRF) modelling uses the co-fluctuations of voxels in V1 with those in other cortical areas to estimate the equivalent of pRFs for the latter (Bock et al., 2015; Gravel et al., 2014; Haak et al., 2013). Estimated cRFs correspond well with pRFs obtained with standard techniques. Voxel-to-voxel multivariate regressions (Baldassano et al., 2012) between V1 and target cortical areas have been used to estimate retinotopic mappings in the latter. The estimated maps fit closely with results from classical retinotopic mappings (Baldassano et al., 2016).

Therefore, the neural cooperation underlying perceptual organization could be reflected in co-fluctuations of fMRI activity between V1 sites. This idea was tested recently by Nasr and co-workers (Nasr et al., 2021) in human subjects. Larger correlations were found between V1 regions of interest (ROIs) in opposite hemispheres when these areas mapped parts of the same compared to parts of different objects. This effect was stronger for stimuli in the lower visual field, congruent with a similar bias for global perception in behavioral data (Nasr & Tootell, 2020; Previc, 1990). These results suggest that changes in perceptual organization are associated with modifications of functional connectivity within V1.

Here we follow up on the Nasr et al. study, using different stimuli and a finer anatomical scale. Global vs. Local compound (i.e., Navon) letters were used (Figure 1A), which induced attention to different spatial scales. We hypothesized that intra-V1 network topology would depend on the perceptual organization of observed scenes. Vertexwise correlation of fMRI activity (between all cortical V1 nodes) was measured in time-windows corresponding to each stimulus. Furthermore, a novel pipeline for intra-V1 connectivity analysis is proposed (Figure 2) to identify reliable differences in network topology across conditions. Both edge-based and network-based tests were employed (Chung et al., 2021; Noble et al., 2022). We mapped links that differed consistently between stimulus classes onto the VF by exploiting the close correspondence between V1 anatomy and retinotopy (Benson et al., 2012; Benson & Winawer, 2018).

**Figure 1.**
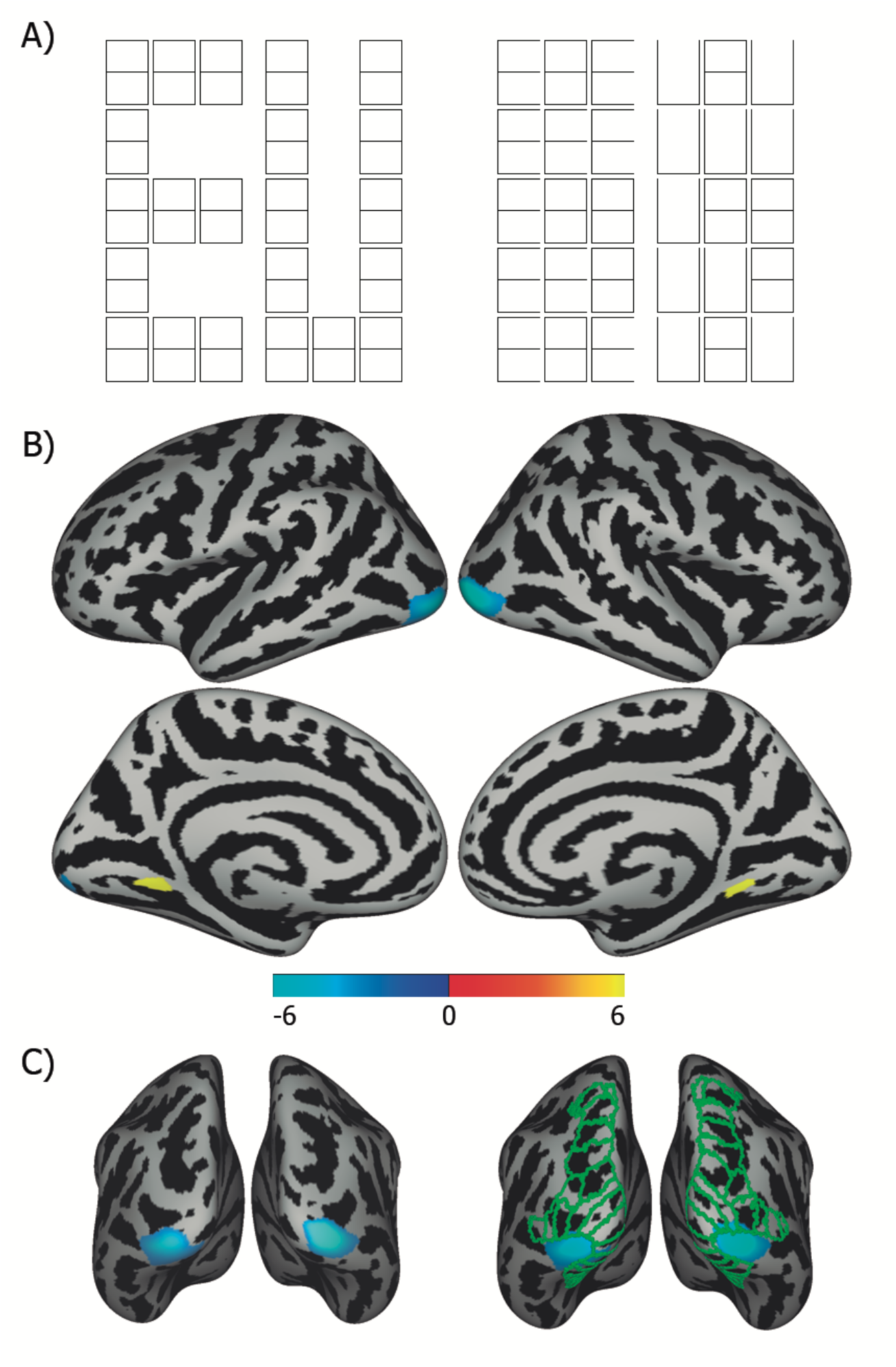
Stimuli and Global>Local activation maps. A) Stimuli used in the experiment. From left to right, Global ‘E’ (EG) and ‘U’ (UG), Local ‘E’ (EL) and ‘U’ (UL). Black/white are reversed. Second-level activation maps of our data (FDR threshold q=0.05), in B) lateral and medial views and in C) posterior views. On the right in C), green lines show the borders of retinotopic areas from the Wang et al. (2015) atlas. The colour bar corresponds to t-values.

**Figure 2.**
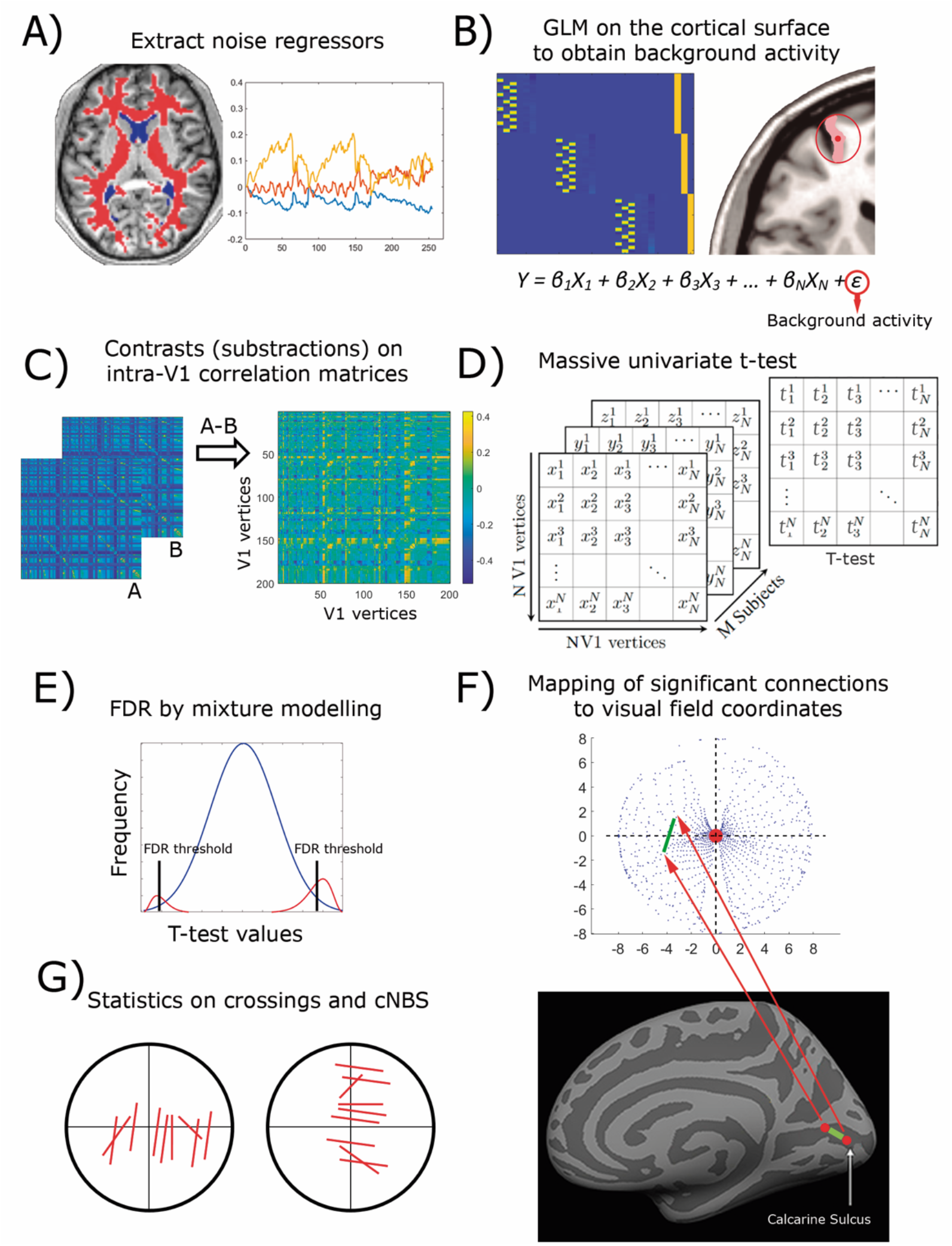
Primary data analysis pipeline. A) The first five principal components of white matter and ventricular (red and blue respectively in the image) time series (and derived measures) were extracted from the volume fMRI data. B) A general linear model on cortical surface data was used to regress out the contributions of these noise parameters, motion correction parameters (and derived measures), and stimulus-evoked activity. C) The resulting background activity was windowed to correspond with stimulus effects (considering the hemodynamic response lag). Correlation matrices between all V1 cortical vertices were obtained per subject, Fisher transformed and subtracted to produce difference matrices. D) A massive univariate t-test was performed on each entry in the difference matrices across subjects. After this step, network-based statistics (cNBS) were calculated in addition to edgewise statistics, as described in the following steps. E) Mixture modelling of the t-values was used to obtain false discovery rate thresholds, illustrated here by a Gaussian null (blue) and two gamma functions (red). F) Connections on the cortex that survived the FDR thresholds were mapped to visual field coordinates (see red arrows), using population values for the pRF centers of each V1 cortical vertex (blue dots). An example of a between-vertex link is shown in green on the inflated cortical surface and the visual field. G) In the edgewise analysis, descriptors of the connections were measured.

We first verified if the intraV1-networks identified here presented three previously reported network properties. One is the dependence of correlations between sites in V1 on their separation along the cortical surface (Dawson et al., 2016). The second property is a reduction of correlations during tasks compared to rest (Ito et al., 2020). The third is that statistical dimensionality increases during tasks compared to rest for brain-wide networks (Ito et al., 2020), reflecting less information redundancy. We then tested our central hypothesis with an edgewise analysis predicting the lengths and spatial distribution of V1-links would correspond with the respective perceptual windows of Global and Local letters (Han et al., 1999). Furthermore, it was predicted that asymmetries in the VF mapping of significant links and perceptual abilities would mirror each other (Barbot et al., 2021; Kupers et al., 2019). Specifically, we expected stronger interhemispheric connections in the lower visual field for Global compared to Local stimuli (consistent with Nasr et al., 2021) and more VF links for Local compared to Global stimuli (see Robertson & Ivry, 2000).

Our final question concerned the relative involvement of different fMRI sources in network generation. The V1 networks could be driven by feedforward responses shared across participants or activity idiosyncratic to each participant. We modelled task fMRI activity as a sum of components representing different sources (Nastase et al., 2019), the weight of which could be modified in the fMRI signal by appropriate preprocessing. This procedure allowed us to assess the role of these sources in intra-V1 connectivity by using a network-based statistical approach (Noble & Scheinost, 2020) while also confirming findings from the edgewise method.

## Results

The intra-V1-network topologies associated with Global/Local Navon letters were compared. We used modified Navon figures that allowed temporal separation of the two levels, thus unmixing their fMRI responses. These stimuli (alternating with background masks) were presented in 20 s blocks, preceded by a Base mask during 20 s, within an MRI scanner for twenty-six participants. Local and Global versions of ‘E’ and ‘U’ were used, and the task was to count minor -and infrequent-deviations in letter shape (see Valdés-Sosa et al., 2020).

A second-level analysis (FDR corrected, q=0.05) found increased activity in the foveal V1 for Local -compared to Global-letters (Figures 1B and C). Previous work has explained this localization by a more restricted spatial focus of attention for local attention (Sasaki et al., 2001). Increased activity was also found for Global -compared to Local-letters in a small region of the para-hippocampal cortex.

### V1 network analysis pipeline

We developed the analysis pipeline depicted in Figure 2 to estimate V1 connectivity based on fMRI activity and segregated by stimuli. In each participant, the fMRI time series were projected from their native volume to their cortical surface and subsequently registered to the FreeSurfer average surface. Nuisance variables were regressed out of the fMRI time series to eliminate the effects of noise, artifacts, and physiological contaminants (Ciric et al., 2017). The primary analysis was carried out on background activity, defined as the residual time series after regressing out the stimulus-evoked activity. The stimulus-evoked activity was modelled by convolving the SPM canonical hemodynamic response function (HRF) with the timing of each stimulus separately. As described in another section, additional variants of Step B from Figure 2, no-deconvolution and deconvolution with finite impulse responses (FIR), were also tested and used to explore the neural sources of V1 networks. Time segments related to the same condition were identified (after accounting for the HRF delay) and concatenated. The fMRI time series from 863 vertices from the V1 area mapping of the central 8° of the visual field as defined in (Benson & Winawer, 2018) were extracted. The Pearson correlations between these time series were calculated.

The second-level connectivity analysis consisted of contrasts between the matrices from different stimulus conditions. Subtracting correlations minimizes the effect of between-site geodesic and retinotopic-eccentricity separation, which otherwise would dominate V1 connectivity estimates. Two within-subject statistical approaches were used. The first consisted of massive univariate t-tests on the connection (edge) strengths, with correction for multiple testing through false discovery rate (FDR, q=0.05) thresholds. Edgewise analysis has the advantage of greater spatial precision but also less statistical power (Noble et al., 2022).

Descriptors related to topology were derived from the networks sparsified with the FDR thresholds, using ten geometric definitions of connectivity subspaces or connSubsp (e.g., links crossing the vertical meridian in the lower visual field, see Figure 7 for cartoon depictions).

We used constrained network-based statistics -cNBS- (Noble and Scheinost, 2020), which is somewhat less precise spatially than edgewise analysis but is more powerful. This method inputs the edgewise tests, which are grouped into “a priori” subnetworks, for which summary statistics are generated and submitted to permutation tests. Here the connSubsp partition was used for cNBS.

### Plausibility check of V1-connectivity matrices

The compatibility of our intra-V1 networks with previous findings in the literature was tested. First, intra-V1 correlations between sites decrease with their separation on the surface (Dawson et al., 2016). As seen in Figure 3 A, the same result was obtained for intra-V1 correlations. Power functions (similar across conditions, y=3.25*x^-0.75^ with R2=0.85 for the Local, and y=3.23*x^-0.74^ with R2=0.82 for the Global letters) explained a large proportion of the relationship between correlations and geodesic distance. This finding validates our strategy of subtracting measures between conditions to eliminate this effect, which would swamp any other influence on the correlations.

**Figure 3.**
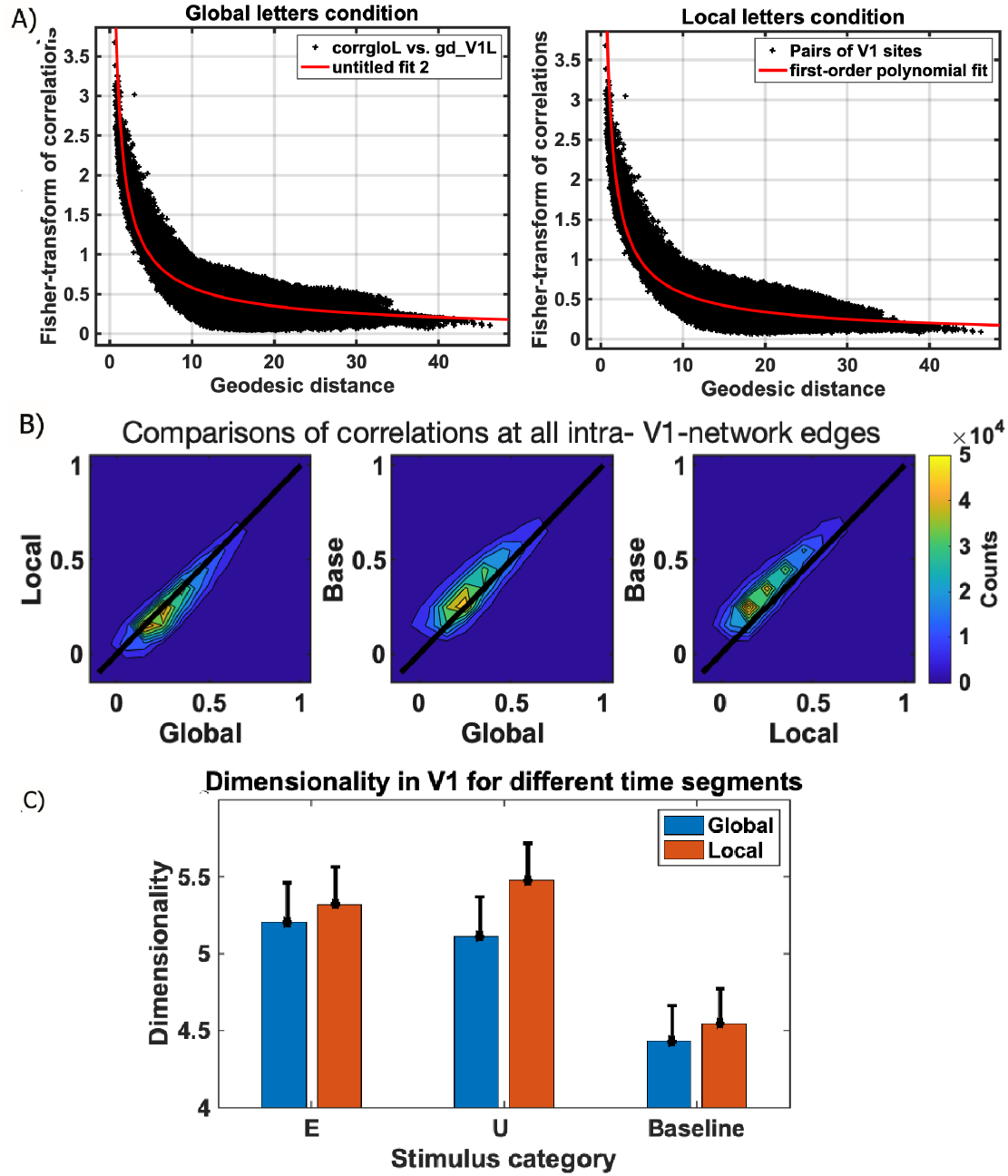
Intra-V1 connectivity plausibility check results. A) Between-site correlations (black dots) in the left V1 as a function of their geodesic separation over the cortical surface. A first-order polynomial regression fit to the data is shown as a red curve for each condition. B) The 2D histograms of binned correlations for pairs of stimuli across all edges in V1. The black diagonal line indicates identical correlation values. Base is the pre-stimulus baseline. C) Dimensionality for different V1-covariance matrices.

Second, correlations in interregional and intra-V1functional networks are smaller for task fMRI than in the resting state (Ito et al., 2020). This reduction was also valid here for our intra-V1 connectivity (Figure 3B). Correlations obtained associated with the baseline (our substitute for resting state) were larger than for the Global and Local segments at 76% and 70 % of the edges, respectively. These effects were significant in paired t-tests on the z-transformed V1 correlations (t(349029)=271.3, p<0.00001 for Global, and t(349029)=429.5, p<0.00001 for Local letters). There was also a significant (t(349029)=,271, p<0.0001), albeit less pronounced, advantage for Global over Local letters (in 24% of edges). Measures of Cohen’s d confirmed this pattern. The effect size was moderate (0.72) for the baseline>Global, strong (1.6) for baseline>Local, and small 0.45) for Global>Local comparisons.

Third, the statistical dimensionality of brain-wide networks increases during tasks compared to the resting state. The V1-covariance matrix dimensionality associated with each stimulus and preceding Base segments (segregated by the level of letters following them) are shown in Figure 3C. Larger dimensionality was found for Global (F(2, 50)=13.0,p=0.0013) and Local (F(2, 50)=16.5, p=0.0004) letters compared to both types of Base. Effects related to letter identity, letter level, or their interaction were absent. Dimensionality for Global relative to Local was also reduced, a non-significant effect that should be verified in a properly powered study. Therefore, our intra-V1 networks exhibited network properties consistent with those reported previously.

### Edgewise analysis of intra-V1 networks associated with Global/Local Letters

Our main goal was to test if V1 networks associated with Global and Local stimuli differed in topology. The results of the edgewise connectivity analyses (with canonical-HRF deconvolution) are shown in Figure 4. FDR thresholds were found at both tails of the distribution of t-values (Figure 4A), probably because distinct network topologies were present for each condition. The link maps in the VF (Figure 4B) show these topological differences. It is evident that for the Global condition, numerous links connected the left and right visual hemifields, especially for the lower visual quadrants. These horizontal –left/right-links were absent for the Local condition, for which most edges remained within one quadrant, especially the inferior right quadrant. Power is reduced in edgewise analyses due to the correction for multiple comparisons. Figure 5 displays links with Cohen’s d values above 0.5. This figure confirms the results of the t-tests and suggests that some links with moderate/large effect sizes did not survive the FDR threshold (which tends to be conservative). FDR threshold estimation failed for the edgewise analyses with the no-deconvolution, FIR deconvolution, and intersubject correlation schemes.

**Figure 4.**
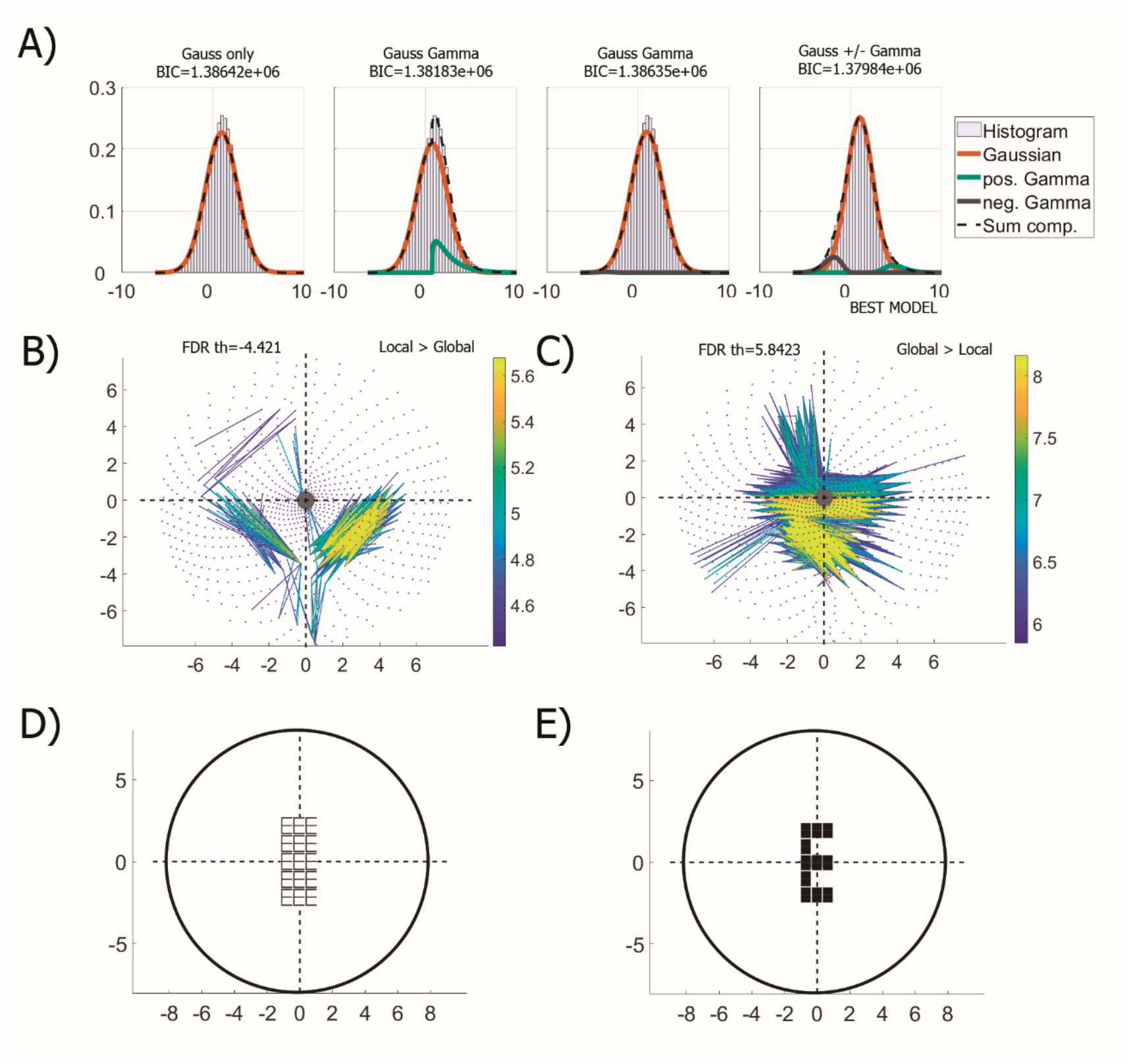
A) Distribution of t-values and mixture models fits. Bayesian information criterion (BIC) values are displayed above each mixture model. B) Plot in the visual field of connections surviving the FDR threshold (q=0.05) for Local>Global contrast, and C) for the Global>Local. Blue dots are pRFs centers for each V1 vertex. FDR threshold in insets. The lines connect the two centers for each significant link, with a color code indicating their absolute t-value. D and E) Approximate size of stimuli in the visual field. As shown in the method section, a narrow sector surrounding the lower half of the vertical meridian had missing data.

**Figure 5.**
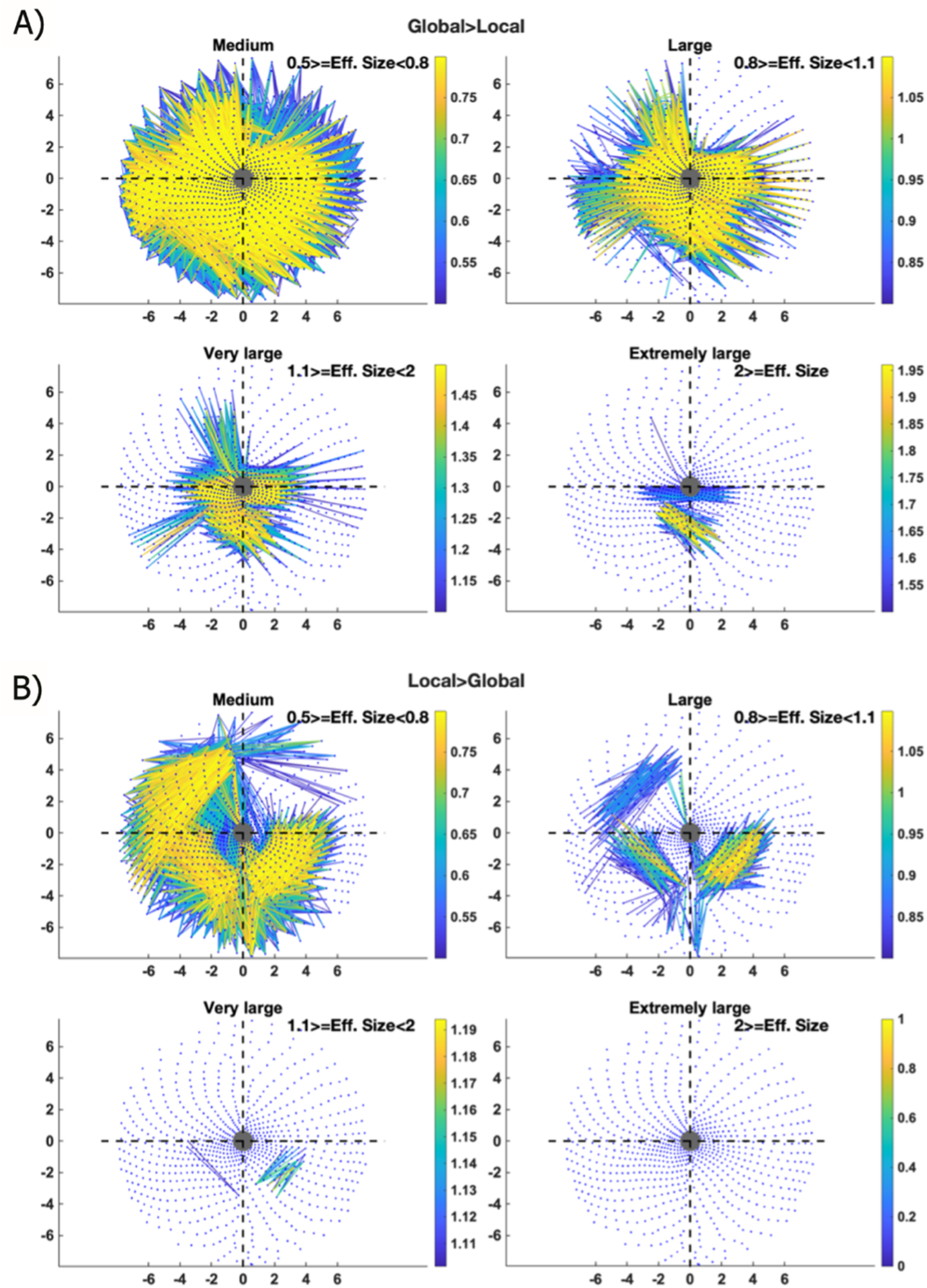
Effect sizes (Cohen’s d) for the V1 links for each contrast (no vertex-wise FDR thresholding). The maps are sorted according to effect size ranges indicated in the insets. Only d > 0.5 are shown. A) Global>local. B) Local>Global.

As predicted, significant connections tended to be longer in visual field space for the Global>Local contrast (median=4.9°) than for the Local>Global contrast (median=3.6°) (Figure 6). This difference in distributions was highly significant in a Kruskal-Wallis test (df=1, Chi-sq=3.6471e03, p<0.0001). Note that the t-values are larger also for the Global compared to the Local conditions. Furthermore, the median length for both types of stimuli was greater than 3° of the visual field, which is the maximum pRF size in central 8° of the visual field (Wandell & Winawer, 2015) studied here. To conclude, the edgewise analysis upheld our main prediction that intra-V1 network topologies were adjusted to the perceptual windows required by Local and by Global stimuli, with links that did not fit within single V1 pRFs.

**Figure 6.**
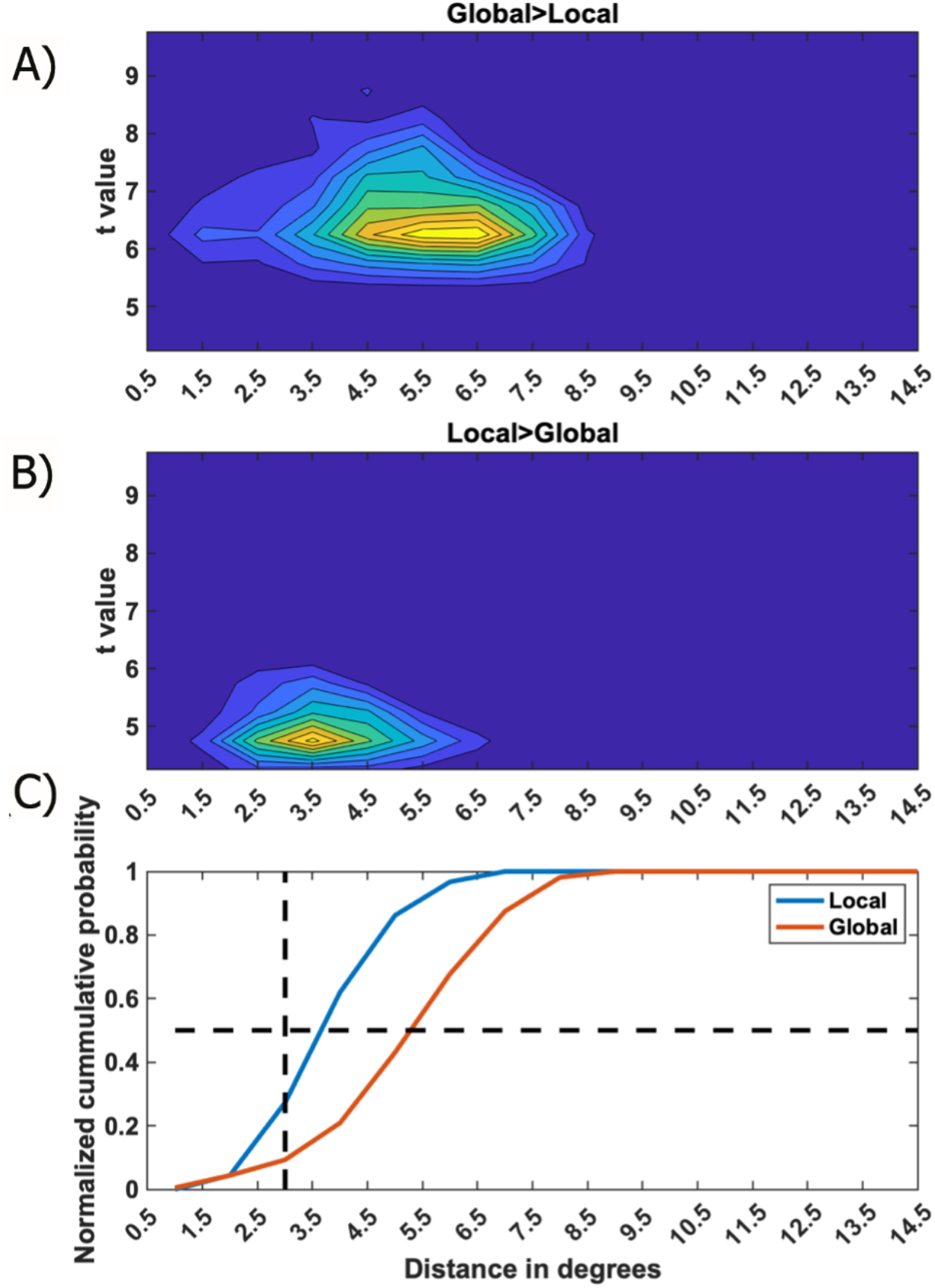
Histograms of link t-values and lengths in the visual field. A) 2-D histogram of link lengths and t value counts for elements surviving the Global>Local FDR threshold; B) 2-D histogram of link lengths and t value counts for elements surviving the Local>Global FDR threshold. C) Normalized cumulative histogram of distances for links in A) and B). The vertical black dashed line corresponds to maximum pRF sizes in V1 for the central 8 degrees of eccentricity. The horizontal line corresponds to the 50% cumulative distribution level (medians).

**Figure 7.**
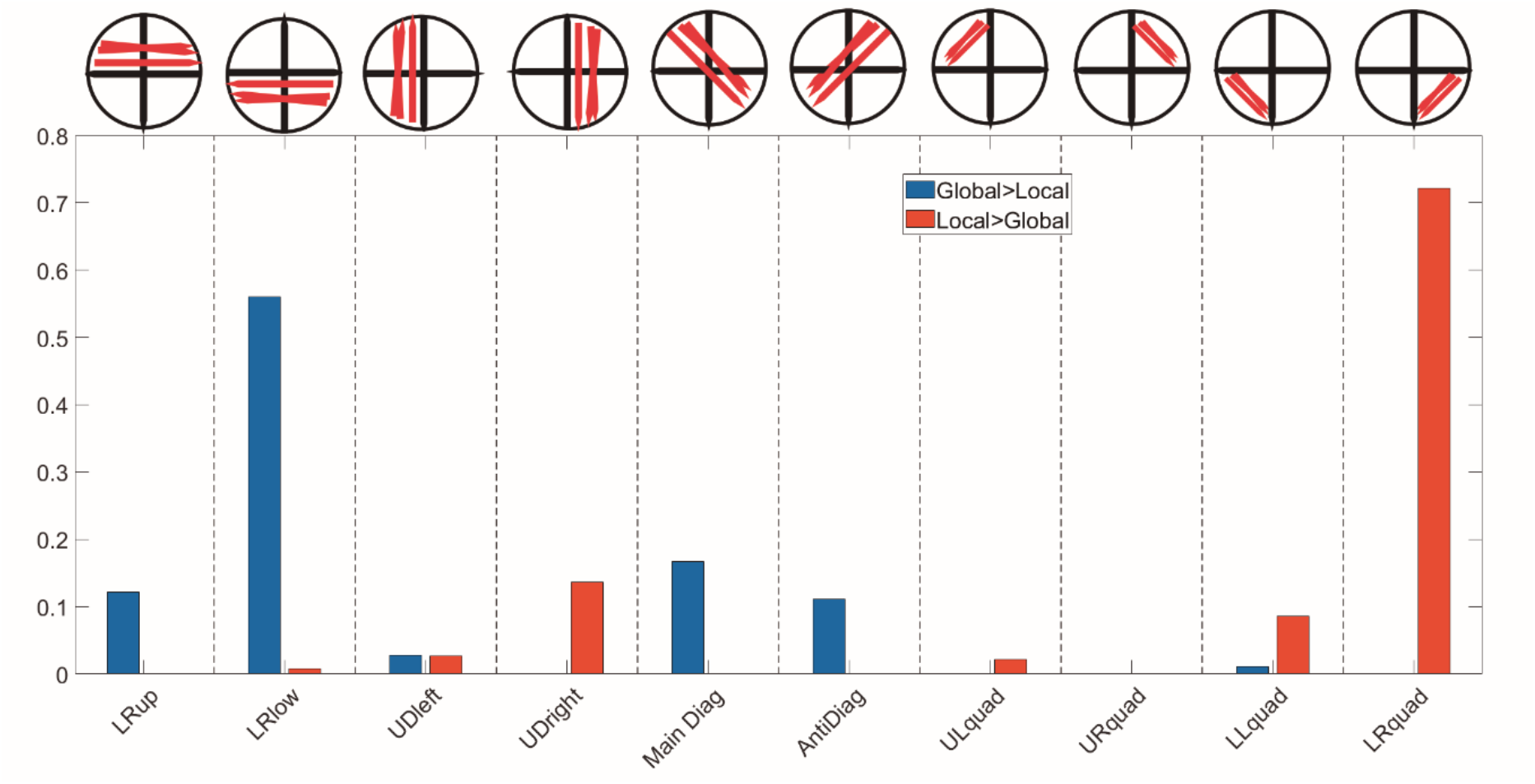
The proportion of connections between/within visual quadrants. Insets schematize the types of connections (connSubsp groups, also used in the network-based analysis). LRup and LRlow, left-right crossings in the upper and lower hemifields. UDleft and UDright, up-down crossings in the left and right hemifields. Main Diag and Antidiag are diagonal links. URquad, ULquad, LLquad, and LRquad, are the links limited to the upper-right, upper-left, lower-left and lower-right visual quadrants.

### Visual field asymmetries in intra-V1 networks

We examined the strength and VF distribution of intra-V1 links that were significant in the Global vs. Local contrasts by measuring the proportion that crossed from one visual quadrant to another or stayed in the same quadrant (Figure 7). For this, we used the connSubsp subnetwork partition. Connections for the Global>Local case mainly crossed the vertical meridian, linking the left and right visual hemifields (i.e., V1 from the two cerebral hemispheres), mainly in the lower quadrants. Diagonal and antidiagonal connections were also present for this condition.

In the Local>Global case, few connections crossed the vertical meridian, and most links stayed within the lower right quadrant (with a smaller number in the lower left quadrant). In this case, links between the upper and lower quadrants on the right side of the VF, something not seen for the Global>Local contrast.

Using Fisher exact tests, we found significant differences between the Global and Local conditions in the proportion of crossings for the following subspaces: lower visual field (p<0.0001), up-down crossings in the left hemifield (p<0.023), up-down in the right (p<0.0001), crossings staying within the lower left (p<0.0001) and lower right quadrants (p<0.0001). Thus, there were evident asymmetries -exclusive for each condition-in link distribution across the VF. As will be discussed below, these effects parallel psychophysical asymmetries.

### Enhancement and suppression of fMRI sources

Different fMRI sources could drive intra-V1 networks, whose relative contributions can be selectively modulated at the GLM step in our pipeline. The stimulus sequence and timing were identical for all participants. Hence, it was possible to model the fMRI signal at each cortical vertex as the sum of four sources, according to the following equation (modified from Nastase et al., 2019):

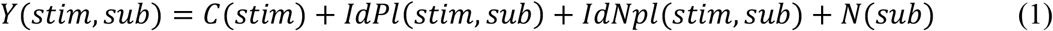

Where *stim* is the stimulus, *sub* is the participant, *Y* is the fMRI time series, *C* is the response shared across participants, *IdPl* is the idiosyncratic phase-locked response to the stimuli, *IdNpl* is the idiosyncratic non-phase-locked response, and *N* represents noise components. Since *N* was reduced in all analyses by regressing out nuisance covariates, the used signals would mainly contain mixtures of the first three terms of equation 1. Table 1 shows how different fMRI preprocessing schemes can theoretically suppress specific components.

**Table 1.** Effects of preprocessing on fMRI sources for intra-V networks.

| Components contributing to correlations after each preprocessing strategy |  |  |  |
| --- | --- | --- | --- |
| | $C(stim)$ | $IdPl(stim, sub)$ | $IdNpl(stim, sub)$ |
| ISC | Keep | - | - |
| No-deconvolution | Keep | Keep | Keep |
| Deconvolve with Canonical HRF | - | Keep | Keep |
| Deconvolve with FIR | - | - | Keep |

Intersubject correlations (ISCs) were estimated between the average fMRI time series at each V1 vertex across n-1 participants and the time series of the other vertices from the left-out participant (these correlation matrices were subsequently averaged). Averaging will isolate the shared response *C*. In the no-deconvolution scheme, task-events responses were not regressed out (preserving *C, IdPl a*nd *IdNp*). Deconvolving task effects with the canonical HRF (as used in previous studies, Keller et al., 2022; Tran et al., 2018) suppresses *C*, leaving *IdPl and dNpl*. Here the canonical HRF captured *C* variance well, with significant correlations in V1 sites (0.8-1.0) between the grand averages and the time course modelled with this HRF. Deconvolving responses to task-events with an FIR basis suppresses the phase-locked components (Goutte et al., 2000; Kay et al., 2008), leaving only *IdNpl*.

Our characterization of the effects of preprocessing schemes (Table 1) is sustained by the power spectra of the resulting fMRI time series (Figure 8). Base + stimulus blocks lasted 44 s implying a fundamental frequency of 0.023 Hz. The Grand average (dominated by the *C* component) presents prominent peaks at this frequency, its harmonics, and the main subharmonic (probably generated by signal aliasing). These peaks are smaller in the raw data, lesser after canonical HRF deconvolution, and smallest after FIR deconvolving.

**Figure 8.**
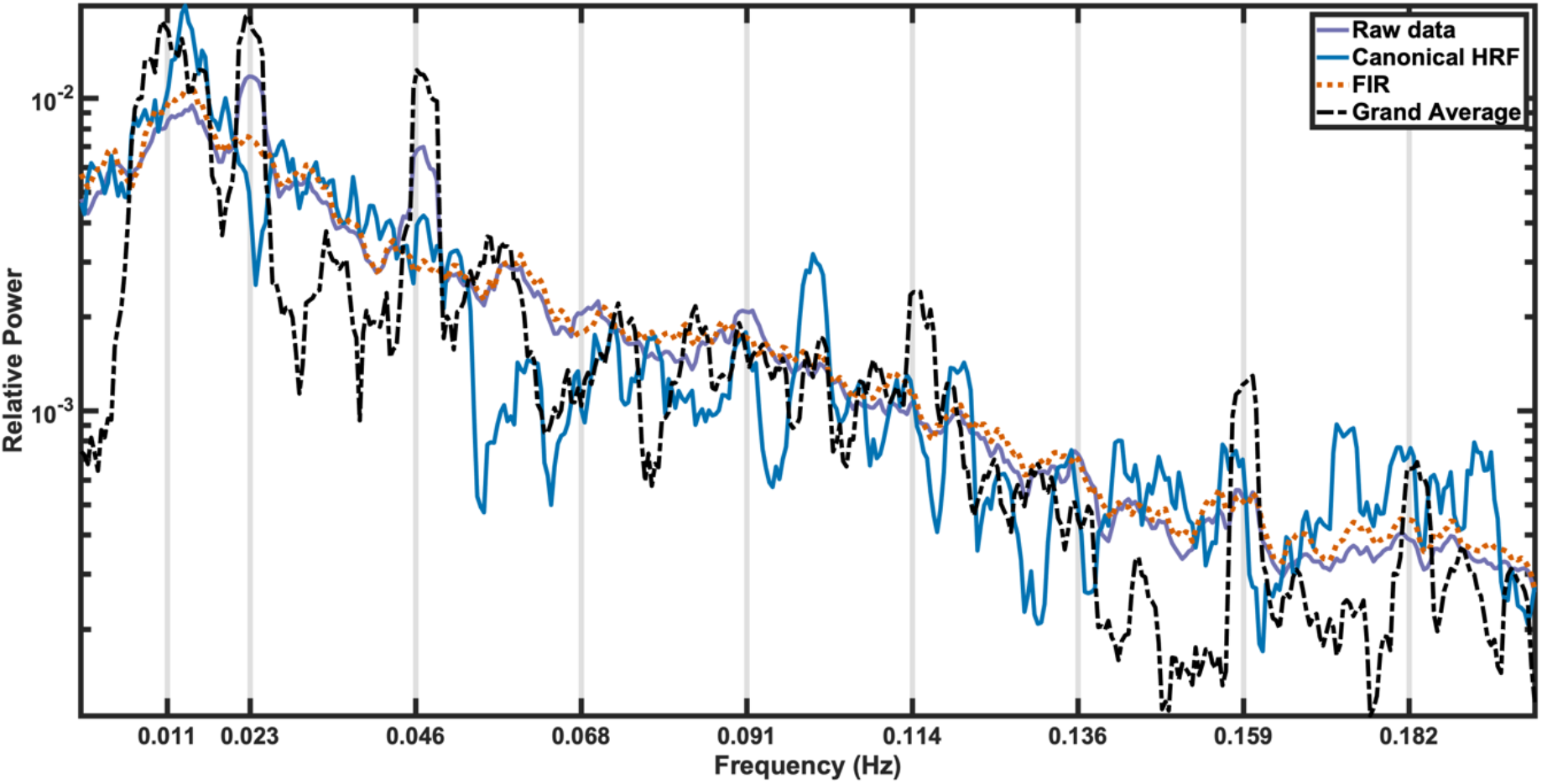
Relative fMRI power spectra. Each curve represents results from a different preprocessing strategy. All spectra were calculated after denoising. The vertical gray lines represent the fundamental frequency of stimulation (1/44 s), the first seven harmonics, and the 0.5 sub-harmonic. The grand average is the spectrum of the time series after averaging across subjects. Raw is the average of spectra across participants when the stimulus-response was not deconvolved. Canonical HRF and FIR are the averages of spectra across participants after the stimulus-response was regressed out using the respective basis functions.

### Network-based analysis (cNBS)

We confirmed results from the edge-based analysis with a constrained network-based statistic (cNBS), which has greater statistical power (Noble et al., 2022). We employed the “a priori” connSubsp partition to avoid circularity. In this analysis, we also compared the effects of different preprocessing schemes. The cNBS in each connSubsp group was defined as the mean t-values of its links Global Vs. Local contrast (restricted to the vertices in the central 4° of eccentricity). A null distribution for the cNBS of each subnetwork was created by randomly flipping the signs of the Global Vs. Local contrast across participants and then recalculating the cNBS 10,000 times. The probability of the observed cNBS in the null distribution was Bonferroni corrected for multiple tests (shown in Table 2).

**Table 2.**
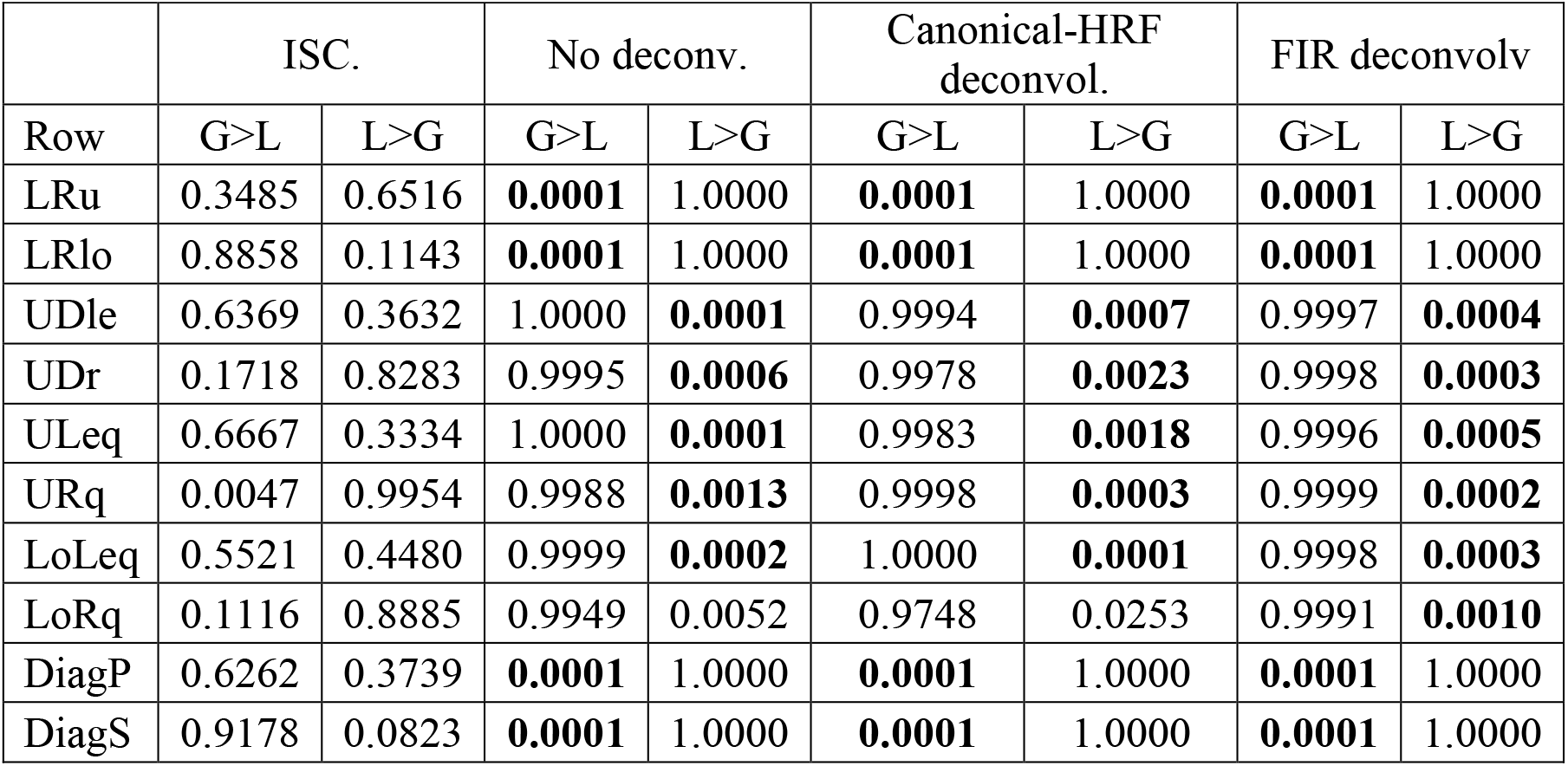
cNBS permutation test results as a function of preprocessing type. Entries are the probability of belonging to the null distribution. With Bonferroni correction, p=0.05 is 0.005, significant results in bold font.

Results from the cNBS analysis are consistent with those from the edgewise tests for the time series deconvolved with the classic HRF (compare with Figures 5 and 6). Subnetworks crossing the vertical meridian (LRu and LRlo) or that were diagonal (DiagP and DiagS) were significantly above chance for the Global>Local contrast. Conversely, subnetworks crossing the horizontal meridian or those restricted to one quadrant were significantly above chance for the Local>Global contrast. These findings were also valid with no deconvolution or with FIR deconvolution. In contrast, none of the tests using ISC (representing activity common to all subjects) were significant. This analysis was confirmed by bootstrap (across participants) estimates of the cNBS.

The 95% Confidence intervals (CIs) excluded zero for all subnetworks with no-deconvolution, canonical HRF- and FIR-deconvolution. In sharp contrast, all CIs for ISC included zero. There was a trend to larger cNBS in the following order: no-deconvolution, HRFF-deconvolution and FIR-deconvolution. However, this was confirmed by bootstrapping the differences in cNBS between preprocessing variants only with uncorrected p-values (p<0.01) and for the Global>Local contrast. These results demonstrate that the intra-V1 networks described in this article do not originate from the *C* component -common to all participants-relying instead on contributions to the fMRI that are characteristic of each individual.

## Discussion

Intra-V1, like brain-wide networks, presented an overall decrease of correlations during stimulation compared to baseline but with increased statistical dimensionality. Patterns of synchrony in neural activity between V1 cortical nodes varied systematically with re-configurations in perceptual organization. Compared with Local letters, Global Navon letters were associated with longer connections in the visual field that frequently crossed the vertical meridian, especially in the lower visual field. In contrast, links related to Local letters were shorter and restricted mainly to the lower right visual quadrant and only infrequently crossed the vertical meridian. These effects were not seen in the intersubject correlations.

Our intra-V1 networks had characteristics previously reported for brain networks. Firstly, intra-V1 correlations decreased with greater separation between cortical nodes, as reported in other studies (Dawson et al., 2016; Raemaekers et al., 2014). We circumvented this significant effect due to geodesic distance by subtracting the correlation matrices associated with different conditions. Secondly, we found that correlations within V1 dropped with Navon letter stimulation (compared to the Base), whereas statistical dimensionality increased. The same outcome has been reported for brain-wide networks using large datasets (Ito et al., 2020), suggesting less information redundancy between brain regions during tasks, possibly boosting information coding capacity. Finally, the peaks of fMRI power spectra in our V1 data exhibited the typical 1/f slope and changed as predicted with stimulus deconvolution.

The edgewise and network-based methods produced consistent results, with all links that were significant in the first analysis belonging to significant sub-networks of the latter. The edgewise analysis was more conservative, with FDR threshold estimation failing in some analyses. Consequently, edgewise testing, followed by correction for multiple comparisons, only allows visualization of ‘tip of the iceberg’ effects (Chen et al., 2022). This idea is reinforced by the many links with large Cohen d values that did not survive FDR thresholds. However, t-maps (especially with canonical HRF deconvolution) allowed more precise visualization of network topology changes in the VF.

Our results are congruent with Nasr et al., 2021. In both studies, the synchrony of fMRI activity between the left and right V1 sites increased when they mapped parts of the same object compared to parts of different objects. In our case, Global letters straddled the vertical meridian of the VF, requiring inter-hemispheric cooperation to perceive their overall shape. Like the fragmented condition in Nasr et al., most of our local letters were circumscribed to either the left or right visual VFs, precluding the need for interhemispheric integration. The segregation of most local letters to one VF would explain this case’s lack of left/right links. Attention to local letters in compound figures implies perceptual segregation into smaller elements (Han & Humphreys, 2002), consistent with the short V1 links confined to single quadrants.

Moreover, in both studies, the significant interhemispheric connections were more robust in dorsal V1 (i.e., the lower VF). Visual perception is thought to be more Global in the lower than the upper VF (Christman, 1993; Levine & McAnany, 2005; Previc, 1990), perhaps related to greater sensitivity for lower spatial frequency (SF) components in this region compared to the upper VF. Lower SF components are critical for extracting Global shapes in Navon figures, whereas higher SF components are more important for perceiving Local shapes (Flevaris & Robertson, 2016). The neural basis of this vertical asymmetry has been recently examined (Nasr & Tootell, 2020).

Links related to Local letters were mainly restricted to the lower right VF (projecting to the left hemisphere). When different Navon letters are briefly flashed to each VF, this competition favours right-sided stimuli when attending the local level and prefers left-sided stimuli when attending the global level(Flevaris et al., 2010). Patients with left hemisphere lesions perform worse on recognizing Local Navon letters than those with right hemisphere lesions (Robertson & Ivry, 2000), who fail more on Global letters. Furthermore, attention to Local Navon letters elicits larger event-related potentials over the left side of the scalp (Han et al., 2002; Iglesias-Fuster et al., 2014; Jiang & Han, 2005), whereas Global Navon letters elicit larger responses over the right side. The left/right cerebral hemispheres are thought to be specialized for Local/Global processing (Flevaris & Robertson, 2016; Han & Humphreys, 2002), an idea which fits the concentration of links associated with Local letters in the right VF.

In contrast to Nasr and co-workers, we found for the Local>Global contrast significant correlations between the dorsal and ventral V1 (corresponding to lower/upper VFs) from the same hemisphere. These connections crossing the horizontal meridian perhaps signify augmented vertical neural binding. The reasons for this discrepancy are unclear, but we list two hypotheses to explore. First, we used a more extensive set of nuisance covariates (as standard in functional connectivity work, e.g., Cole et al., 2021) than Nasr et al., which could have increased sensitivity by lowering noise. Second, we used actively attended foveal stimuli, whereas unattended peripheral stimuli were used in Nasr et al. The former approach could perhaps strengthen links crossing the horizontal meridian.

The feedforward response to the stimuli (*C*) shared among participants (Nastase et al., 2019) does not contribute to the intra-V1 networks described here. Although the leave-one-subject-out procedure used to calculate ISC can inflate positive findings (Chen et al., 2016), we found only null results with this method. Conversely, the networks are present after deconvolution with the canonical HRF or FIR (procedures that suppress *C*). These findings implicate fMRI activity idiosyncratic to the participants as the network source. Idiosyncratic activity can be phase locked to the stimuli (e.g., fluctuations in evoked activity amplitude or duration) or non-phased locked (e.g., spontaneous oscillations, also known as background activity).

A second clue to the neural sources of the networks is the lengths of significant links in the VF, which exceed the largest pRF sizes in the V1 sectors mapping our stimuli (Benson & Winawer, 2018; Wandell & Winawer, 2015). Connection lengths implicate either long-range lateral interactions within V1 (Kapadia et al., 1995; Liang et al., 2017) or feedback from higher-order visual areas with large RFs (Briggs, 2020; Williams et al., 2008; Petro & Muckli, 2017). It is thought (Field & Hayes, 2004; Kuai et al., 2017; Schwarz & Bolz, 1991) that enhanced co-activation of neurons via lateral connections in the superficial layers of V1 can facilitate the detection of extended contours. Microelectrode recordings in monkeys suggest that contour detection emerges first in V4 and then is fed back to V1 (Chen et al., 2014; Liang et al., 2017). Layer-specific fMRI studies in humans have found effects of global stimulus configuration in the superficial layers of V1 (Lawrence et al., 2019; Marquardt et al., 2019; Petro & Muckli, 2016), possibly originating from feedback originating in other cortical areas.

Joint fMRI and electrophysiological recordings are required to understand the V1 networks described here. Ultra-slow spontaneous fMRI fluctuations are reported to be associated with the envelope of gamma-band neural activity (Nir et al., 2007; Scheeringa et al., 2011; Mateo et al., 2017). Gamma-band activity could affect vasomotor oscillations in cortical surface pial arterioles, which in turn would influence the arrival of oxyhemoglobin to cortical tissue, consequently changing the fMRI signal (Mateo et al., 2017; Drew et al., 2020). A seminal hypothesis (Gray et al., 1989) is that perceptual binding of contours is produced by the synchronous neuronal activity of neurons representing different aspects of the same object (Singer, 2021), although not all agree (Riesenhuber & Poggio, 1999). Neuronal synchronization in beta-band neural activity has also been proposed as a player in contour integration (Kermani et al., 2020).

One limitation of this study is the lack of eye-movement control, which would allow stimuli to stray in the VF, blurring the V1 retinotopic mapping. Another limitation is that we used a population prior map (Benson & Winawer, 2018) to plot V1 vertices onto the VF when individualized retinotopic mapping would be more precise. To register cortical anatomy across participants, we used the default FreeSurfer inter-subject surface registration (Fischl, 2012) instead of more accurate methods (e.g., MSMSULC and MSMALL, Coalson et al., 2018). Due to missing values, the significance of link effects in a narrow sector of the VF may be underestimated. A caveat for follow-up studies is that our methods in their present form are optimal for block designs for which clear temporal separation between experimental conditions is guaranteed. Furthermore, it would be wise only to compare networks associated with stimuli differing minimally in retinal layout.

### Ideas and speculations

V1 networks, estimated from background fMRI activity, seem to reflect the perceptual organization of observed visual scenes, with stable topologies across individuals. Although the spatial resolution of these networks is coarse compared to microelectrode recordings in monkeys (and blind to faster neuronal interactions), they speak to a finer scale than ROI-based networks. Furthermore, they are easy to estimate in human experiments, and their visualization in the VF benefits from the close correspondence between retinotopy and anatomy found in V1. They could occupy an intermediate level of analysis of V1 circuitry, perhaps analogous to fMRI-based pRFs relative to microelectrode-measured RFs. Several testable predictions of this idea immediately come to mind.

Intra-V1 networks could facilitate object recognition in noisy backgrounds, help circumvent camouflage, and guide object-based attention, with success in these tasks co-varying alongside network efficiency. The operation of Gestalt laws and different interpretations of ambiguous (Wyatte et al., 2014), or multi-stable figures (Li et al., 2017), may be reflected in V1 networks. Their topology may also vary as attention shifts within the same stimuli (Flevaris et al., 2011; Luck et al., 1997). Canonical and non-canonical orientations stimulus orientations (e.g., upright vs. inverted faces, Strother et al., 2011) could also be associated with different V1 network configurations. Successful and aborted induction of illusory surfaces (e.g., Kanisza figures) (Chen et al., 2020; Keane et al., 2021) could also generate different network topologies.

The physiological basis of intra-V1 networks could be confirmed in ultrahigh-field fMRI studies with cortical-layer specific recordings (Huber et al., 2020; Marquardt et al., 2019; Petro & Muckli, 2017). Together with granger causality, or mediation analysis, layer-specific recordings could be used to dissect the roles of lateral interaction in V1 from feedback influence from higher-order areas in the association of network topology with perceptual organization.

### Conclusions

Correlations in intra-V1 networks are decreased during stimulation compared to baseline for most links, with specific increases in some connections. The result is larger statistical dimensionality. Thus, perception organization could sculpt V1 connectivity. Global and Local letters elicited different V1 topologies that partially mirror VF perceptual asymmetries. Time-locked responses shared across participants do not contribute to these networks, whose link lengths in the VF frequently exceeded V1 pRF sizes. These networks possibly reflect interactions within V1 or feedback to this area from higher-order areas, which would modulate fMRI activity idiosyncratic to each participant. These findings, and further studies with the methods we developed, could help shed light on V1 as a “cognitive blackboard”. The main appeal of studying intra-V1 networks is that it may allow indirect but precise visualization of cooperative effects among higher-order visual areas, otherwise hidden in voxels scattered across the cortex.

## Materials and Methods

### Participants

Twenty-six human volunteers aged from 23 to 28 years (9 females) participated in the study. All had normal, or corrected-to-normal, vision did not present any medical condition and were right-handed except for two cases. The ethics committee of the University for Electronic Science and Technology of China (UESTC) approved the procedures, and participants gave written informed consent following the Helsinki declaration.

### Task and stimuli

This experiment is described in more detail in Valdés-Sosa et al., 2020. The stimuli were projected on a screen at the subject’s feet, viewed through an angled mirror fixed to the MRI head-coil, and were generated using the Cogent Matlab toolbox (http://www.vislab.ucl.ac.uk/cogent.php). Modified Global and Local Navon letters, made from white lines on a black background (Figure 1A), alternated with a baseline matrix of (about 2.0° wide and 5.3° high). This matrix was built out of smaller placeholder elements shaped like ‘8’s (with visual angles about 40’ wide and 1° 3’ high). Two letters, ‘E’ and ‘U’, were presented at both levels. Blocks consisted of ten 1 sec repetitions of the same letter (within a fixed level) that alternated with 1 sec baseline periods, thus lasting 20 s. Blocks were initiated by a 1 s cue (‘Global’ or ‘Local’), followed by a 19 s baseline, and ended with a 4 s wait period where counts of minor deviations in letter shape were reported. Four or five runs were presented, each consisting of 12 blocks.

### Data acquisition and basic image preprocessing

Data acquisition and initial image preprocessing are described in detail in Valdés-Sosa et al., 2020. Recordings were carried out with a GE Discovery MR750 3T scanner (General Electric Medical Systems, Milwaukee, WI, USA) using an eight-channel receiver head coil. A T1-weighted image was obtained with 1 × 1 × 0.5 mm resolution. A mid-gray surface was reconstructed from the T1 image for each subject using Freesurfer (http://surfer.nmr.mgh.harvard.edu), registered to the FsAverage template, and subsampled to 81924 vertices. Discs with 5mm radii were defined over the FsAverage surface using the Surfing toolbox (http://surfing.sourceforget.net). The T1 image was also segmented and normalized to MNI space using SPM12. For each subject, masks of white matter and of cerebrospinal fluid (CSF) restricted to the ventricles (using a template in MNI space) (https://sites.google.com/site/mrilateralventricle/template were created using a threshold of tissue probability greater than 0.9.

Functional images were obtained with a T2*-weighted echo planar imaging sequence (TR=2.5s; TE =40 ms; flip angle=90^○^). Spatial resolution was 1.875 × 1.875 × 2.9, with 135 images per run. The initial five volumes in all runs were discarded to stabilize T1 magnetization. Pre-preprocessing of functional data included artifact correction (ArtRepair toolbox, http://cibsr.stanford.edu/tools/ArtRepair/ArtRepair.htm), followed by slice-timing, head motion correction (with the extraction of motion parameters) and unwarping using SPM8 (http://www.fil.ion.ucl.ac.uk/spm/). The fMRI time series were then projected to each individual’s mid-gray cortical surface and high-pass filtered with a time constant of 128 s.

### Preprocessing of fMRI time series and intra-V1 connectivity estimates

Background activity was defined as the residual time series of each surface vertex after regressing out the effects of the stimuli (evoked response) and 64 nuisance parameters using the general linear model (GLM). The nuisance regressors included the six primary motion parameters (obtained from head motion correction), their derivatives, and the quadratics of both these sets (24 motion regressors in total). Physiologic noise was modelled using the aCompCor method (Behzadi et al., 2007) on the time series extracted separately from the white matter and the ventricles masks described above. The first five principal components from each set of time series, the derivatives of these components, and the quadratics of all these parameters were obtained (40 regressors in total). Therefore, the nuisance regressors totaled 64 parameters. We applied a univariate general linear model (GLM) with the vertex-wise fMRI time series as dependent variables in three variants. The first variant modelled the stimulation block as a square wave convolved with the canonical hemodynamic function from the SPM-12 toolbox (canonical-HRF). In a second variant (no-deconvolve), only nuisance regressors were included in the design matrix. In the last variant, a FIR basis (18 stick functions covering 45 s: the approximate duration of a stimulus cycle) was used. Intersubject correlations were also calculated, as described in the results section. All four variants were used in the network-based statistical analysis. We verified with a Fourier analysis the effects of these preprocessing schemes on the periodic stimulus-related activity. The power spectra of complete time series were extracted for each individual and averaged over participants. We used Bartlett’s method to calculate the spectra, which involves averaging the periodograms of consecutive and non-overlapping segments. Theoretically, this method does not have the best statistical properties. However, after comparisons with multi-taper spectrum estimation, little difference was found in the spectral estimates and much less over-smoothing of peaks with the simpler method.

After applying the GLM, the time series were segmented into blocks considering the time shift introduced by the hemodynamic function. These segments were linearly detrended. Segments corresponding to the same stimulus were concatenated. The activity from surface vertices was smoothed by averaging with the time series of its neighbours in the 5 mm discs mentioned above. For this article, we used only vertices corresponding to the central 8° of eccentricity in a population prior retinotopic map of V1 (Benson & Winawer, 2018). Different vertices could be missing due to noise or BOLD signal dropout. These bad vertices were concentrated in a small subset of the central fovea and a narrow circular sector about 10 degrees wide, surrounding the lower portion of the vertical visual field meridian (see Figure 10). Data from all participants was obtained in 560 V1 vertices, whereas missing data for one case was found in 85 and for two participants in 68 vertices. The maximum number of missing participants was 7 (in 3 vertices). Thus, caution should be exercised in interpreting results from the visual field sectors with the most missing data. Still, the conclusions about the overall differences between connectivity for Global and Local stimuli are not undercut by this fact.

**Figure 9.**
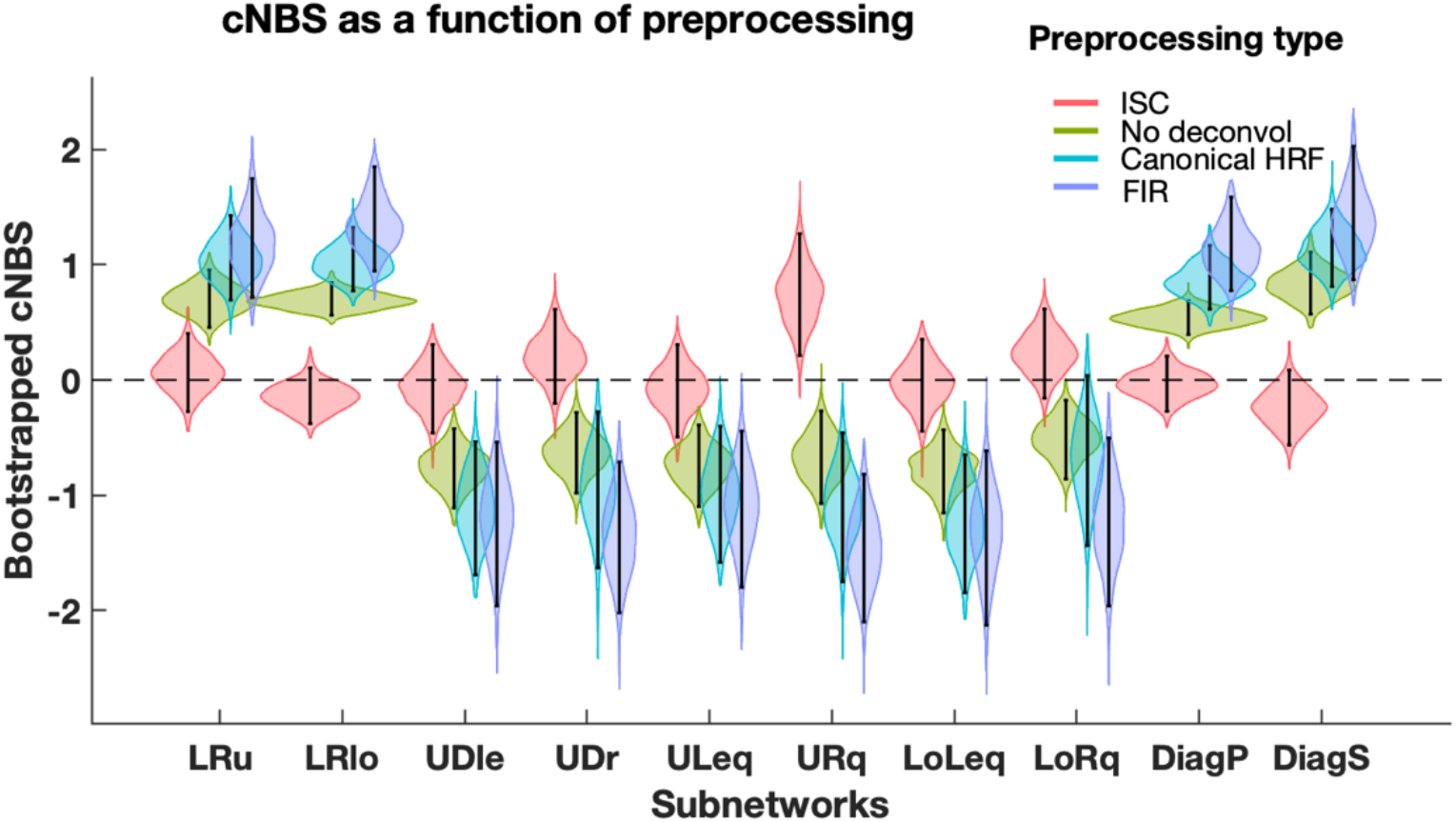
Violin plot of bootstrap cNBS replicates in the Global-Local contrast as a function of preprocessing type. cNBS was calculated for the ten connSubsp by resampling participants with replacement (n=1000). Positive and negative values indicate larger cNBS values for the Global and Local conditions, respectively. Black dots correspond to means, the black line to 99% confidence intervals, and the violin plot colour to preprocessing type.

**Figure 10.**
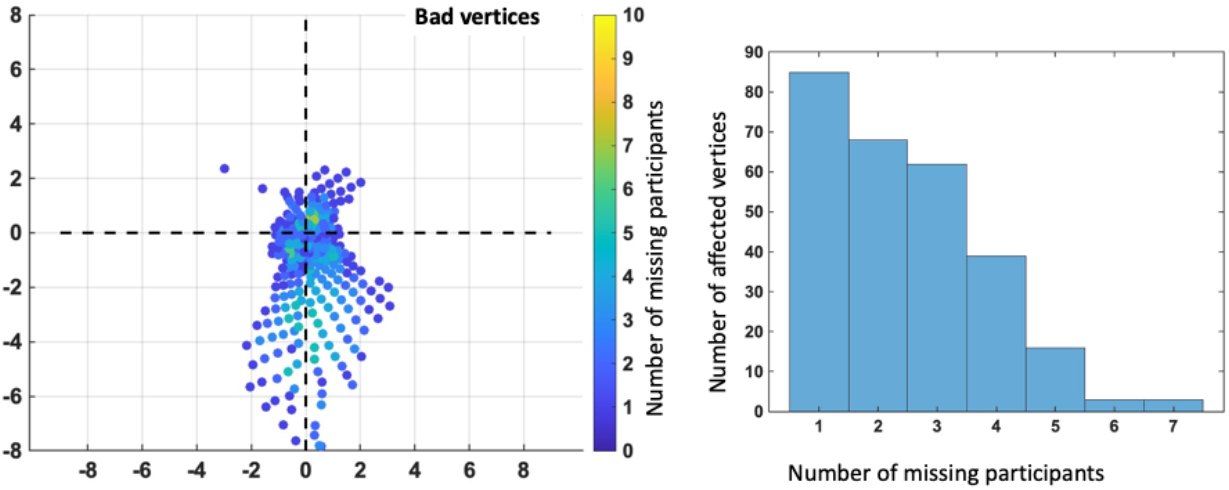
Missing V1 vertices. The left side displays a plot of how many participants had missing data at each measured location within the visual field (axis conventions for this graph are the same as in Figure 1). On the right side, a histogram of how many vertices (y-axis) had missing data as a function of the number of participants in which this happened (x-axis).

Covariance and Pearson correlation matrices in V1 were calculated based on these time series for each participant and stimulus condition and transformed to z values using the Fisher transformation. Negative correlations were set to zero (Sporns & Betzel, 2016), and missing values in each subject’s V1-correlation matrix were substituted by the median value of the other subjects

### Plausibility check of V1-connectivity matrices

As a plausibility check of the V1-connectivity matrices, three previously described properties of fMRI networks were tested. First, we tested the relationship between the correlation and geodesic distance between V1. The geodesic distance between V1 cortical sites for each cortical hemisphere (based on the FsAverage surface) was calculated with the Fast Marching toolbox (https://es.mathworks.com/matlabcentral/fileexchange/6110-toolbox-fast-marching). The Fisher-transformed correlations were fit as power functions of geodesic distance. MATLAB curve-fit toolbox. Second, the size of correlations from resting state (here represented by a pre-stimulation Base period) and task fMRI were compared. Correlations for the two stimulus conditions were also compared. The values from the group average of the V1-correlation matrices for each pair of conditions were used to calculate 2-dimensional histogram density plots. Edgewise differences between conditions were submitted to groupwise t-tests. Finally, the dimensionality of resting state and task fMRI data was also verified. Statistical dimensionality for fMRI data was estimated by the ‘participation ratio’ (Litwin-Kumar et al., 2017), as used by Ito et al. (Ito et al., 2020). The V1-covariance matrices were obtained for each condition and their preceding baseline, and their eigenvalues were extracted. Roughly explained, the participation ratio finds the number of eigenvalues needed to explain variance greater than a given threshold, with more components indicating a higher dimensionality of the data. Dimensionality values were submitted to a one-way repeated-measures analysis of variance across stimulus conditions.

### Edgewise analysis

Within-subjects, two-tailed, group t-tests were calculated based on the differences between Global and Local connectivity matrices. Multiple comparison corrections were based False Discovery Rates (FDR) with q=0.05. FDRs thresholds were estimated for both tails of the t-value distribution by using mixture modelling (Bielczyk et al., 2018; Gorgolewski et al., 2012) that estimated a Null (H0) and alternative distribution (H1s) from the histogram of observed t-values. Four mixture models were fit using expectation-maximization: a) Only a Gaussian-H0; b) The Gaussian H0 plus a negative gamma H1 distribution; c) the Gaussian-H0 plus a positive gamma H1 distribution; d) And the Gaussian-H0 plus negative gamma and positive gamma H1 distributions. The model with the lowest Bayesian Information Criterion (BIC) was the best fit (Gorgolewski et al., 2012). The FDR was calculated for each t-value as the quotient of the area under the H0 distribution above the observed t-value relative to the number of counts of the empirical histogram (which has contributions from H0 and H1). The FDR was calculated for both tails of the histogram. The minimum/maximum t-value in the right/left tails associated with an FDR less than q was defined as the threshold. For some preprocessing variants, this threshold estimation failed.

### Constrained network-based statistics (cNBS)

Ten sub-networks (connSubsp) of the intra-V1 connectivity matrix were predefined (illustrated in the top row of Figure 7). The number of connections falling into each sub-network was counted and displayed in histograms, and Fisher’s exact test was carried out to contrast the conditions. The cNBS methodology (Noble & Scheinost, 2020) was used for statistical tests of the differences between Global and Local conditions. The cNBS was calculated by averaging the edgewise t-values within each connSubsp sub-network. The null distribution of the cNBS in each subnetwork was obtained by permutation resampling (flipping the signs of the observed t-values over participants, n=10,000) and recalculating the statistic. The p-value for the observed cNBS in each group was calculated from the null distribution. Moreover, confidence intervals for the subnetwork cNBSs were obtained by bootstrapping over participants 1000 times.

### Projection of V1 connectivity onto the visual space

The mean polar angle and eccentricity were extracted for V1 cortical vertices from a population prior map (Benson & Winawer, 2018). Graph plots depicted significant edges from the edgewise tests in visual field coordinates. The colour of lines in the graph plot represented either the t-tests surviving the FDR thresholds or Cohen’s d binned according to effect strength. The length of these lines was measured in visual degrees, and the difference in distributions between experimental conditions was assessed with a Kruskal-Wallis test.

## Author contributions

Designed research; MVS, MOO

Contributed analytic tools: MVS, MOO, ALC, PVS Analyzed data: MVS, MOO, JIF

Wrote the paper: MVS, MOO, JIF, PVS

## Acknowledgements

We thank the University of Electronic Science and Technology of China (UESTC) for funding and providing special equipment and facilities for this project. This work was supported by the VLIR-UOS project “A Cuban National School of Neurotechnology for Cognitive Aging”, the National Fund for Science and Innovation of Cuba, the National Natural Science Foundation of China (#81861128001) and the 1110 Project (B12027) of China. The authors are grateful to K. Uludag for his suggestions on the manuscript.

## Bibliography

Baldassano, C., Fei-Fei, L., & Beck, D. M. (2016). Pinpointing the peripheral bias in neural scene processing networks during natural viewing. Journal of Vision, 16(2), 9. https://doi.org/10.1167/16.2.9.doi

Baldassano, C., Iordan, M. C., Beck, D. M., & Fei-Fei, L. (2012). Voxel-level functional connectivity using spatial regularization. NeuroImage, 63(3), 1099–1106. https://doi.org/10.1016/j.neuroimage.2012.07.046

Barbot, A., Xue, S., & Carrasco, M. (2021). Asymmetries in visual acuity around the visual field. Journal of Vision, 21(1), 1–23. https://doi.org/10.1167/JOV.21.1.2

Behzadi, Y., Restom, K., Liau, J., & Liu, T. T. (2007). A component based noise correction method (CompCor) for BOLD and perfusion based fMRI. NeuroImage, 37(1), 90–101. https://doi.org/10.1016/j.neuroimage.2007.04.042

Benson, N. C., Butt, O. H., Datta, R., Radoeva, P. D., Brainard, D. H., & Aguirre, G. K. (2012). The retinotopic organization of striate cortex is well predicted by surface topology. Current Biology : CB, 22(21), 2081–2085. https://doi.org/10.1016/j.cub.2012.09.014

Benson, N. C., & Winawer, J. (2018). Bayesian analysis of retinotopic maps. ELife, 7, 1–29. https://doi.org/10.7554/elife.40224

Bielczyk, N. Z., Walocha, F., Ebel, P. W., Haak, K. v., Llera, A., Buitelaar, J. K., Glennon, J. C., & Beckmann, C. F. (2018). Thresholding functional connectomes by means of mixture modeling. NeuroImage, 171(October 2017), 402–414. https://doi.org/10.1016/j.neuroimage.2018.01.003

Bijsterbosch, J., Harrison, S. J., Jbabdi, S., Woolrich, M., Beckmann, C., Smith, S., & Duff, E. P. (2020). Challenges and future directions for representations of functional brain organization. Nature Neuroscience, 23(12), 1484–1495. https://doi.org/10.1038/s41593-020-00726-z

Bijsterbosch, J., Smith, S., & Beckmann, C. (2017). Introduction to resting state fMRI functional connectivity. Oxford University Press.

Bock, A. S., Binda, P., Benson, N. C., Bridge, H., Watkins, K. E., & Fine, I. (2015). Resting-State Retinotopic Organization in the Absence of Retinal Input and Visual Experience. Journal of Neuroscience, 35(36), 12366–12382. https://doi.org/10.1523/JNEUROSCI.4715-14.2015

Briggs, F. (2020). Role of Feedback Connections in Central Visual Processing. Annual Review of Vision Science, 6, 313–334. https://doi.org/10.1146/annurev-vision-121219-081716

Chen, G., Shin, Y. W., Taylor, P. A., Glen, D. R., Reynolds, R. C., Israel, R. B., & Cox, R. W. (2016). Untangling the relatedness among correlations, part I: Nonparametric approaches to inter-subject correlation analysis at the group level. NeuroImage, 142, 248–259. https://doi.org/10.1016/j.neuroimage.2016.05.023

Chen, G., Taylor, P. A., Stoddard, J., Cox, R. W., Bandettini, P. A., & Pessoa, L. (2022). Sources of Information Waste in Neuroimaging: Mishandling Structures, Thinking Dichotomously, and Over-Reducing Data. Aperture Neuro, 2021(5), 1–22. https://doi.org/10.52294/2e179dbf-5e37-4338-a639-9ceb92b055ea

Chen, M., Yan, Y., Gong, X., Gilbert, C. D., Liang, H., & Li, W. (2014). Incremental Integration of Global Contours through Interplay between Visual Cortical Areas. Neuron, 82(3), 682– 694. https://doi.org/10.1016/j.neuron.2014.03.023

Chen, S., Weidner, R., Zeng, H., Fink, G. R., Müller, H. J., & Conci, M. (2020). Tracking the completion of parts into whole objects: Retinotopic activation in response to illusory figures in the lateral occipital complex. NeuroImage, 207(June 2019), 116426. https://doi.org/10.1016/j.neuroimage.2019.116426

Klink, P.C., Chen, X., Vanduffel, W., & Roelfsema, P. R. (2021). Population receptive fields in non-human primates from whole-brain fmri and large-scale neurophysiology in visual cortex. ELife, 10, 1–35. https://doi.org/10.7554/eLife.67304

Christman, S. D. (1993). Local-global processing in the upper versus lower visual fields. Bulletin of the Psychonomic Society, 31(4), 275–278. https://doi.org/10.3758/BF03334927

Chung, J., Bridgeford, E., Arroyo, J., Pedigo, B. D., Saad-Eldin, A., Gopalakrishnan, V., Xiang, L., Priebe, C. E., & Vogelstein, J. T. (2021). Statistical connectomics. Annual Review of Statistics and Its Application, 8, 463–492. https://doi.org/10.1146/annurev-statistics-042720-023234

Ciric, R., Wolf, D. H., Power, J. D., Roalf, D. R., Baum, G. L., Ruparel, K., Shinohara, R. T., Elliott, M. A., Eickhoff, S. B., Davatzikos, C., Gur, R. C., Gur, R. E., Bassett, D. S., & Satterthwaite, T. D. (2017). Benchmarking of participant-level confound regression strategies for the control of motion artifact in studies of functional connectivity. NeuroImage, 154(March), 174–187. https://doi.org/10.1016/j.neuroimage.2017.03.020

Coalson, T. S., van Essen, D. C., & Glasser, M. F. (2018). The impact of traditional neuroimaging methods on the spatial localization of cortical areas. Proceedings of the National Academy of Sciences of the United States of America, 115(27), E6356–E6365. https://doi.org/10.1073/pnas.1801582115

Cole, M. W., Ito, T., Cocuzza, C., & Sanchez-Romero, R. (2021). The functional relevance of task-state functional connectivity. Journal of Neuroscience, 41(12), 2684–2702. https://doi.org/10.1523/JNEUROSCI.1713-20.2021

Dawson, D. A., Lam, J., Lewis, L. B., Carbonell, F. M., Mendola, J. D., & Shmuel, A. (2016). Partial Correlation-Based Retinotopically Organized Resting-State Functional Connectivity Within and Between Areas of the Visual Cortex Reflects More Than Cortical Distance. Brain Connectivity, 6(1), 57–75. https://doi.org/10.1089/brain.2014.0331

DeValois, R. L., & DeValois, K. K. (1991). Spatial Vision. In Oxford Psychology Series. https://doi.org/10.1093/acprof:oso/9780195066579.001.0001

Drew, P. J., Mateo, C., Turner, K. L., Yu, X., & Kleinfeld, D. (2020). Ultra-slow oscillations in fMRI and resting-state connectivity: neuronal and vascular contributions and technical confounds. Neuron, 107(5), 782–804.

Field, D. J., & Hayes, A. (2004). Contour Integration and the Lateral Connections of V1 Neurons. In The Visual Neurosciences (pp. 1096--1079).

Fischl, B. (2012). FreeSurfer. NeuroImage, 62(2), 774–781. https://doi.org/10.1016/j.neuroimage.2012.01.021

Flevaris, A. v, Bentin, S., & Robertson, L. C. (2010). Local or global? Attentional selection of spatial frequencies binds shapes to hierarchical levels. Psychological Science, 21(3), 424– 431. https://doi.org/10.1177/0956797609359909

Flevaris, A. v., Bentin, S., & Robertson, L. C. (2011). Attentional selection of relative SF mediates global versus local processing: Evidence from EEG. Journal of Vision, 11(7), 1– 12. https://doi.org/10.1167/11.7.1

Flevaris, A. v., & Robertson, L. C. (2016). Spatial frequency selection and integration of global and local information in visual processing: A selective review and tribute to Shlomo Bentin. Neuropsychologia, 83, 192–200. https://doi.org/10.1016/j.neuropsychologia.2015.10.024

Goodale, M. A., & Milner, A. D. (2018). Two visual pathways – Where have they taken us and where will they lead in future? Cortex, 98, 283–292. https://doi.org/10.1016/j.cortex.2017.12.002

Gorgolewski, K. J., Storkey, A. J., Bastin, M. E., & Pernet, C. R. (2012). Adaptive thresholding for reliable topological inference in single subject fMRI analysis. Frontiers in Human Neuroscience, 6(AUGUST), 1–14. https://doi.org/10.3389/fnhum.2012.00245

Goutte, C., Nielsen, F. Å., & Hansen, L. K. (2000). Modeling the haemodynamic response in fMRI using smooth FIR filters. IEEE Transactions on Medical Imaging, 19(12), 1188– 1201. https://doi.org/10.1109/42.897811

Gravel, N., Harvey, B., Nordhjem, B., Haak, K. V., Dumoulin, S. O., Renken, R., Ćurčić-Blake, B., & Cornelissen, F. W. (2014). Cortical connective field estimates from resting state fMRI activity. Frontiers in Neuroscience, 8(OCT), 1–10. https://doi.org/10.3389/fnins.2014.00339

Gray, C. M., König, P., Engel, A. K., & Singer, W. (1989). Oscillatory responses in cat visual cortex exhibit inter-columnar synchronization which reflects global stimulus properties. Nature, 338(6213), 334–337. https://doi.org/10.1038/338334a0

Haak, K. v., & Beckmann, C. F. (2016). Objective analysis of the topological organization of the human cortical visual connectome suggests three visual pathways. Cortex, 1–11. https://doi.org/10.1016/j.cortex.2017.03.020

Haak, K. v., Winawer, J., Harvey, B. M., Renken, R., Dumoulin, S. O., Wandell, B. A., & Cornelissen, F. W. (2013). Connective field modeling. NeuroImage, 66, 376–384. https://doi.org/10.1016/j.neuroimage.2012.10.037

Han, S., & Humphreys, G. W. (2002). Segmentation and selection contribute to local processing in hierarchical analysis. Quarterly Journal of Experimental Psychology Section A: Human Experimental Psychology, 55(1), 5–21. https://doi.org/10.1080/02724980143000127

Han, S., Humphreys, G. W., & Chen, L. (1999). Parallel and competitive processes in hierarchical analysis: perceptual grouping and encoding of closure. Journal of Experimental Psychology. Human Perception and Performance, 25(5), 1411–1432. https://doi.org/10.1037/0096-1523.25.5.1411

Han, S., Weaver, J. A., Murray, S. O., Kang, X., Yund, E. W., & Woods, D. L. (2002). Hemispheric asymmetry in global/local processing: Effects of stimulus position and spatial frequency. NeuroImage, 17(3), 1290–1299. https://doi.org/10.1006/nimg.2002.1255

Harrison, S. A., & Tong, F. (2009). Decoding reveals the contents of visual working memory in early visual areas. Nature, 458(7238), 632–635. https://doi.org/10.1038/nature07832

Huber, L., Finn, E. S., Chai, Y., Goebel, R., Stirnberg, R., Stöcker, T., Marrett, S., Uludag, K., Kim, S. G., Han, S. H., Bandettini, P. A., & Poser, B. A. (2020). Layer-dependent functional connectivity methods. Progress in Neurobiology, May, 101835. https://doi.org/10.1016/j.pneurobio.2020.101835

Iglesias-Fuster, J., Santos-Rodríguez, Y., Trujillo-Barreto, N., & Valdés-Sosa, M. J. (2014). Asynchronous presentation of global and local information reveals effects of attention on brain electrical activity specific to each level. Frontiers in Psychology, 5(OCT), 1–14. https://doi.org/10.3389/fpsyg.2014.01570

Ito, T., Brincat, S. L., Siegel, M., Mill, R. D., He, B. J., Miller, E. K., Rotstein, H. G., & Cole, M. W. (2020). Task-evoked activity quenches neural correlations and variability across cortical areas. PLoS Computational Biology, 16(8 August), e1007983. https://doi.org/10.1371/JOURNAL.PCBI.1007983

Jiang, Y., & Han, S. (2005). Neural mechanisms of global/local processing of bilateral visual inputs: an ERP study. Clinical Neurophysiology : Official Journal of the International Federation of Clinical Neurophysiology, 116(6), 1444–1454. https://doi.org/10.1016/j.clinph.2005.02.014

Kapadia, M. K., Ito, M., Gilbert, C. D., & Westheimer, G. (1995). Improvement in visual sensitivity by changes in local context: Parallel studies in human observers and in V1 of alert monkeys. Neuron, 15(4), 843–856. https://doi.org/10.1016/0896-6273(95)90175-2

Kay, K. N., David, S. V., Prenger, R. J., Hansen, K. A., & Gallant, J. L. (2008). Modeling low-frequency fluctuation and hemodynamic response timecourse in event-related fMRI. Human Brain Mapping, 29(2), 142–156. https://doi.org/10.1002/hbm.20379

Keane, B. P., Barch, D. M., Mill, R. D., Silverstein, S. M., Krekelberg, B., & Cole, M. W. (2021). Brain network mechanisms of visual shape completion. NeuroImage, 236. https://doi.org/10.1016/j.neuroimage.2021.118069

Keller, A. S., Jagadeesh, A. v., Bugatus, L., Williams, L. M., & Grill-Spector, K. (2022). Attention enhances category representations across the brain with strengthened residual correlations to ventral temporal cortex. NeuroImage, 249(January), 118900. https://doi.org/10.1016/j.neuroimage.2022.118900

Kermani, M., Zavitz, E., Oakley, B., Price, N. S. C., Hagan, M. A., & Wong, Y. T. (2020). Longrange neural coherence encodes stimulus information in primate visual cortex. BioRxiv. https://doi.org/10.1101/2020.06.22.164269

Kimchi, R., Yeshurun, Y., Spehar, B., & Pirkner, Y. (2016). Perceptual organization, visual attention, and objecthood. Vision Research, 126, 34–51. https://doi.org/10.1016/j.visres.2015.07.008

Kuai, S. G., Li, W., Yu, C., & Kourtzi, Z. (2017). Contour Integration over Time: Psychophysical and fMRI Evidence. Cerebral Cortex, 27(5), 3042–3051. https://doi.org/10.1093/cercor/bhw147

Kupers, E. R., Carrasco, M., & Winawer, J. (2019). Modeling visual performance differences ‘around’ the visual field: A computational observer approach. PLoS Computational Biology, 15(5), 1–29. https://doi.org/10.1371/journal.pcbi.1007063

Lamme, V. A. F., & Roelfsema, P. R. (2000). The distinct modes of vision offered by feedforward and recurrent processing. Trends in Neurosciences, 23(11), 571–579. https://doi.org/10.1016/S0166-2236(00)01657-X

Lawrence, S. J. D., Formisano, E., Muckli, L., & de Lange, F. P. (2019). Laminar fMRI: Applications for cognitive neuroscience. NeuroImage, 197(March), 785–791. https://doi.org/10.1016/j.neuroimage.2017.07.004

Levine, M. W., & McAnany, J. J. (2005). The relative capabilities of the upper and lower visual hemifields. Vision Research, 45(21), 2820–2830. https://doi.org/10.1016/j.visres.2005.04.001

Li, H.-H., Rankin, J., Rinzel, J., Carrasco, M., & Heeger, D. J. (2017). Attention model of binocular rivalry. Proceedings of the National Academy of Sciences, 114(30), E6192– E6201. https://doi.org/10.1073/pnas.1620475114

Liang, H., Gong, X., Chen, M., Yan, Y., Li, W., & Gilbert, C. D. (2017). Interactions between feedback and lateral connections in the primary visual cortex. Proceedings of the National Academy of Sciences of the United States of America, 114(32), 8637–8642. https://doi.org/10.1073/pnas.1706183114

Litwin-Kumar, A., Harris, K. D., Axel, R., Sompolinsky, H., & Abbott, L. F. (2017). Optimal Degrees of Synaptic Connectivity. Neuron, 93(5), 1153-1164.e7. https://doi.org/10.1016/j.neuron.2017.01.030

Luck, S. J., Chelazzi, L., Hillyard, S. a, & Desimone, R. (1997). Neural mechanisms of spatial selective attention in areas V1, V2, and V4 of macaque visual cortex. Journal of Neurophysiology, 77(1), 24–42.

Markov, N. T., Misery, P., Falchier, A., Lamy, C., Vezoli, J., Quilodran, R., Gariel, M. A., Giroud, P., Ercsey-Ravasz, M., Pilaz, L. J., Huissoud, C., Barone, P., Dehay, C., Toroczkai, Z., Van Essen, D. C., Kennedy, H., & Knoblauch, K. (2011). Weight consistency specifies regularities of macaque cortical networks. Cerebral Cortex, 21(6), 1254–1272. https://doi.org/10.1093/cercor/bhq201

Markov, N. T., Vezoli, J., Chameau, P., Falchier, A., Quilodran, R., Huissoud, C., Lamy, C., Misery, P., Giroud, P., Ullman, S., Barone, P., Dehay, C., Knoblauch, K., & Kennedy, H. (2014). Anatomy of hierarchy: Feedforward and feedback pathways in macaque visual cortex. Journal of Comparative Neurology, 522(1), 225–259. https://doi.org/10.1002/cne.23458

Marquardt, I., de Weerd, P., Schneider, M., Gulban, O. F., Ivanov, D., & Uludağ, K. (2019). Depth-resolved ultra-high field fMRI reveals feedback contributions to surface motion perception. BioRxiv, 1–28. https://doi.org/10.1101/653626

Mateo, C., Knutsen, P.M., Tsai, P.S., Shih, A.Y. & Kleinfeld, D. (2017). Entrainment of arteriole vasomotor fluctuations by neural activity is a basis of blood-oxygenation-level-dependent “resting state” connectivity. Neuron, 96, 1–13.

Nasr, S., Kleinfeld, D., & Polimeni, J. R. (2021). The Global Configuration of Visual Stimuli Alters Co-Fluctuations of Cross-Hemispheric Human Brain Activity. The Journal of Neuroscience, 41(47), 9756–9766. https://doi.org/10.1523/jneurosci.3214-20.2021

Nasr, S., & Tootell, R. B. H. (2020). Asymmetries in global perception are represented in nearversus far-preferring clusters in human visual cortex. Journal of Neuroscience, 40(2), 355– 368. https://doi.org/10.1523/JNEUROSCI.2124-19.2019

Nastase, S. A., Gazzola, V., Hasson, U., & Keysers, C. (2019). Measuring shared responses across subjects using intersubject correlation. Social Cognitive and Affective Neuroscience, 14(6), 669–687. https://doi.org/10.1093/scan/nsz037

Nir, Y., Fisch, L., Mukamel, R., Gelbard-Sagiv, H., Arieli, A., Fried, I., & Malach, R. (2007). Coupling between neuronal firing rate, gamma LFP, and BOLD fMRI is related to interneuronal correlations. Current biology, 17(15), 1275–1285.

Noble, S., Mejia, A. F., Zalesky, A., & Scheinost, D. (2022). Improving power in functional magnetic resonance imaging by moving beyond cluster-level inference. Proceedings of the National Academy of Sciences, 119(32), e2203020119. https://doi.org/10.1073/PNAS.2203020119

Noble, S., & Scheinost, D. (2020). The Constrained Network-Based Statistic: A New Level of Inference for Neuroimaging. In Lecture Notes in Computer Science (including subseries Lecture Notes in Artificial Intelligence and Lecture Notes in Bioinformatics): Vol. 12267 LNCS (pp. 458–468). https://doi.org/10.1007/978-3-030-59728-3_45

Petro, L. S., & Muckli, L. (2016). The brain’s predictive prowess revealed in primary visual cortex. Proceedings of the National Academy of Sciences, 113(5), 1124–1125. https://doi.org/10.1073/pnas.1523834113

Petro, L. S., & Muckli, L. (2017). The laminar integration of sensory inputs with feedback signals in human cortex. Brain and Cognition, 112, 54–57. https://doi.org/10.1016/j.bandc.2016.06.007

Pomerantz, J. R., & Portillo, M. C. (2015). Emergent Features, Gestalts, and Feature Integration Theory. In J. Wolfe & L. C. Robertson (Eds.), From Perception to Consciousness. Oxford University Press. https://doi.org/10.1093/acprof:osobl/9780199734337.003.0016

Previc, F. H. (1990). Functional specialization in the lower and upper visual fields in humans: Its ecological origins and neurophysiological implications. Behavioral and Brain Sciences, 13(3), 519–542. https://doi.org/10.1017/S0140525X00080018

Raemaekers, M., Schellekens, W., van Wezel, R. J. A., Petridou, N., Kristo, G., & Ramsey, N. F. (2014). Patterns of resting state connectivity in human primary visual cortical areas: A 7T fMRI study. NeuroImage, 84, 911–921. https://doi.org/10.1016/j.neuroimage.2013.09.060

Riesenhuber, M., & Poggio, T. (1999). Are cortical models really bound by the “binding problem”? Neuron, 24(1), 87–93. https://doi.org/10.1016/S0896-6273(00)80824-7

Robertson, L. C., & Ivry, R. (2000). Hemispheric asymmetries: Attention to visual and auditory primitives. Current Directions in Psychological Science, 9(2), 59–63. https://doi.org/10.1111/1467-8721.00061

Roelfsema, P. R., & de Lange, F. P. (2016). Early Visual Cortex as a Multiscale Cognitive Blackboard. Annual Review of Vision Science, 2(1), 131–151. https://doi.org/10.1146/annurev-vision-111815-114443

Sasaki, Y., Hadjikhani, N., Fischl, B., Liu, A. K., Marret, S., Dale, A. M., & Tootell, R. B. H. (2001). Local and global attention are mapped retinotopically in human occipital cortex. Proceedings of the National Academy of Sciences, 98(4), 2077–2082. https://doi.org/10.1073/pnas.98.4.2077

Scheeringa, R., Fries, P., Petersson, K. M., Oostenveld, R., Grothe, I., Norris, D. G., … & Bastiaansen, M. C. (2011). Neuronal dynamics underlying high-and low-frequency EEG oscillations contribute independently to the human BOLD signal. Neuron, 69(3), 572–583.

Schwarz, C., & Bolz, J. (1991). Functional specificity of a long-range horizontal connection in cat visual cortex: A cross-correlation study. Journal of Neuroscience, 11(10), 2995–3007. https://doi.org/10.1523/jneurosci.11-10-02995.1991

Singer, W. (2021). Recurrent dynamics in the cerebral cortex: Integration of sensory evidence with stored knowledge. Proceedings of the National Academy of Sciences, 118(33). https://doi.org/10.1073/pnas.2101043118

Smith, F. W., & Muckli, L. (2010). Nonstimulated early visual areas carry information about surrounding context. Proceedings of the National Academy of Sciences of the United States of America, 107(46), 20099–20103. https://doi.org/10.1073/pnas.1000233107

Sporns, O., & Betzel, R. F. (2016). Modular brain networks. Annual Review of Psychology, 67(September 2015), 613–640. https://doi.org/10.1146/annurev-psych-122414-033634

Strother, L., Mathuranath, P. S., Aldcroft, A., Lavell, C., Goodale, M. A., & Vilis, T. (2011). Face inversion reduces the persistence of global form and its neural correlates. PLoS ONE, 6(4). https://doi.org/10.1371/journal.pone.0018705

Tran, S. M., McGregor, K. M., James, G. A., Gopinath, K., Krishnamurthy, V., Krishnamurthy, L. C., & Crosson, B. (2018). Task-residual functional connectivity of language and attention networks. Brain and Cognition, 122(January), 52–58. https://doi.org/10.1016/j.bandc.2018.02.003

Valdés-Sosa, M., Ontivero-Ortega, M., Iglesias-Fuster, J., Lage-Castellanos, A., Gong, J., Luo, C., Castro-Laguardia, A. M., Bobes, M. A., Marinazzo, D., & Yao, D. (2020). Objects seen as scenes: Neural circuitry for attending whole or parts. NeuroImage, 210(January). https://doi.org/10.1016/j.neuroimage.2020.116526

van Essen, D. C., Newsome, W. T., & Maunsell, J. H. R. (1984). The visual field representation in striate cortex of the macaque monkey: Asymmetries, anisotropies, and individual variability. Vision Research, 24(5), 429–448. https://doi.org/10.1016/0042-6989(84)90041-5

Wagemans, J., Feldman, J., Gepshtein, S., Kimchi, R., Pomerantz, J. R., van der Helm, P. a, & van Leeuwen, C. (2012). A century of Gestalt psychology in visual perception: II. Conceptual and theoretical foundations. Psychological Bulletin, 138(6), 1218–1252. https://doi.org/10.1037/a0029334

Wandell, B. A., & Winawer, J. (2015). Computational neuroimaging and population receptive fields. Trends in Cognitive Sciences, 19(6), 349–357. https://doi.org/10.1016/j.tics.2015.03.009

Williams, M. A., Baker, C. I., op de Beeck, H. P., Mok Shim, W., Dang, S., Triantafyllou, C., & Kanwisher, N. (2008). Feedback of visual object information to foveal retinotopic cortex. Nature Neuroscience, 11(12), 1439–1445. https://doi.org/10.1038/nn.2218

Wyatte, D., Jilk, D. J., & O’Reilly, R. C. (2014). Early recurrent feedback facilitates visual object recognition under challenging conditions. Frontiers in Psychology, 5(JUL), 1–10. https://doi.org/10.3389/fpsyg.2014.00674

